# Lessons from Japan marine stock enhancement and sea ranching programmes over 100 years

**DOI:** 10.1101/828798

**Authors:** Shuichi Kitada

## Abstract

More than 26 billion juveniles of 180 marine species are released annually into the wild in over 20 countries, but the usefulness of this strategy remains unclear. Here, I analyse the effects of stocking by Japanese marine and salmon stock-enhancement programmes and evaluate their efficacy through a Bayesian meta-analysis of new and previously considered cases. The posterior mean recapture rate (± SD) was 8.3 ± 4.7%. Without considering personnel costs and negative impacts on wild populations, the mean economic efficiency was 2.8 ± 6.1, with many cases having values of 1 to 2. On the macro-scale, the proportion of released seeds to total catch was 76 ± 20% for Japanese scallop, 28 ± 10% for abalone, 20 ± 5% for swimming crab, 13 ± 5% for kuruma prawn, 11 ± 4% for Japanese flounder, and 7 ± 2% for red sea bream; according to these percentages, stocking effects were generally small, and population dynamics were unaffected by releases but dependent on the carrying capacity of the nursery habitat. All cases of Japanese releases, except for Japanese scallop, were probably economically unprofitable. Captive breeding reduces the fitness of hatchery fish in the wild. In addition, long-term releases replace wild genes and may cause fitness decline in the recipient population when the proportion of hatchery fish is very high. Short-term hatchery stocking can be useful, particularly for conservation purposes, but large-scale programmes may harm the sustainability of populations. Nursery habitat recovery and fishing pressure reduction often outperform hatcheries in the long run.

## Introduction

The history of stocking fish larvae dates back to the 1870s (Blaxter 2000; Svåsand et al. 2000; Bell et al. 2005). Approximately 150 years later, seed production technologies has progressed, and the release of hatchery-reared animals into the wild is now a popular fisheries, forestry, and wildlife management tool (Laikre et al. 2010; Taylor et al. 2017). More than 26 billion juveniles of 180 marine species, including salmonids, are released into the wild every year in more than 20 countries (Kitada 2018). In coastal marine fisheries, uses of hatchery releases are classified into several categories with different objectives: stock enhancement (“stocking cultured organisms to replenish or increase abundance of wild stocks”), sea ranching (“stocking for put-grow-and-take food fisheries”; Leber et al. 2004) and restocking (“restoring depleted spawning biomass to a level of producing regular yields”; Bell et al. 2008). Moreover, marine ranching (basically, hatchery releases accompanied with artificial reef construction in Japan) is rapidly growing in China (Lee & Zhang 2018; Wang et al. 2018; Zhou et al. 2019) and Korea (Lee & Rahman 2018). Despite the huge number of seeds released every year, most studies have been conducted only at the experimental stage (Taylor et al. 2017). In a previous study evaluating the economic performance of 14 cases involving 12 species worldwide, most cases were found to be economically unprofitable because of the high cost of seed production compared with prevailing market prices (Kitada 2018). Empirical studies are still too limited, however, to determine the full effectiveness of hatchery-release efforts (Laikre et al. 2010), and basic questions, namely, “Do hatcheries produce extra fish for harvest, or do they simply replace natural fish with hatchery fish?” and “Are hatcheries cost-effective for producing fish?” have seldom been answered (Waples 1999). More empirical studies are thus obviously needed to settle the controversy of whether hatchery stocking is useful or harmful (Hilborn 1992; Blaxter 2000; Svåsand et al. 2000; Naish et al. 2007; Araki & Schmid 2010; Laikre et al. 2010).

The feasibility and risks of hatchery stocking cannot be tested solely by examining pilot-scale releases—full-scale releases must be considered (Hilborn 2004). Despite this principle, most marine stocking programmes worldwide are in pilot-scales and test enhancement scenarios and release strategies (Taylor et al. 2017). Among the world’s countries, Japan is exceptional, having released the largest number of marine and salmonid species, often on a large scale. Focused and in-depth analyses of Japanese full-scale releases can thus provide one of the best information for improving understanding of the positive and negative effects of marine stock enhancement and sea ranching programmes.

In regard to Japanese marine stock enhancement and sea ranching programmes, several reviews have focused on the effects of stocking several fishes (Masuda & Tsukamoto 1998; Kitada 1999; Kitada & Kishino 2006), kuruma prawn (Hamasaki & Kitada 2006, 2013), abalone (Hamasaki & Kitada 2008a), and decapod crustaceans (Hamasaki & Kitada 2008b; Hamasaki et al. 2011). Attention has also been directed towards seed production (Fushimi 2001; Takeuchi 2001; Le Vay et al. 2007), seed quality and behaviour (Tsukamoto et al. 1999), broodstock management (Taniguchi 2003, 2004), and genetic effects on wild populations (Kitada et al. 2009). Reviews on salmonids have covered large-scale hatchery releases, mainly focusing on chum salmon and pink salmon; however, their emphasis was mainly on ecology, with stocking effects not fully evaluated (e.g., Kaeriyama 1999; Morita et al. 2006; Kaeriyama et al. 2012; Nagata et al. 2012; Miyakoshi et al. 2013; Kitada 2014; Morita 2014). My previous systematic review provided a perspective on economic, ecological, and genetic effects of marine stock enhancement and sea ranching programmes worldwide, including salmon (Kitada 2018), but its focus was global. No integrated review evaluating the results of both marine and salmonid stock enhancement in Japan has yet appeared.

Here, I first outline the history and present status of Japanese salmon and marine hatchery programmes. Second, I evaluate the stocking effects of major full-scale projects using a novel Bayesian meta-analysis, with new cases added to those of the previous study. Third, I predict changes in the contribution of hatchery releases to commercial catches of representative species on a macro-scale. Finally, I summarize the consequences of the world’s largest hatchery stocking programmes with three different types of broodstock management, typically used worldwide: (I) wild collection of larvae, (II) wild collection of parents (may include individuals of hatchery-origin), and (III) captive breeding (Kitada 2018)—namely, those of red sea bream in Kagoshima Bay (Type III, run for 45 years), Japanese scallop (Type I, 50 years), and chum salmon (Type II, more than 100 years). The results obtained here should benefit future fisheries management and conservation practices worldwide.

## Marine and salmonid enhancement programmes in Japan

### Hatcheries and seed release

Japan’s marine stock-enhancement programme was initiated by the Fishery Agency in 1963. At that time, Japan aimed to become a fully developed country under its national industrialization policy. Many Japanese coastlines were reclaimed, mainly up until the 1970s (Fig. 1), and separate coastal industrial zones were created; thereafter, many coastal fishers moved to cities to earn higher incomes, and coastal fisheries failed to grow (Kitada 2001). The marine stock-enhancement programme was a mitigation policy designed to improve degraded habitats and thereby enhance coastal fisheries. Salmon hatcheries have a much longer history in Japan; they were started in the 1880s to increase fishery production of returning chum salmon and have continued for over 130 years (Miyakoshi et al. 2013). As shown later, however, chum salmon stock enhancement failed during the early 1960s. Japan’s marine stock-enhancement programme was therefore started with no successful case precedent. This situation was exactly the same as that described by Hilborn (1992): “Many believe that the future of fisheries lies with artificial propagation. Indeed, throughout the world most management agencies seem to be relying on some form of artificial propagation to rebuild fish stocks that are depleted due to poor fisheries management or poor habitat management.”

**Figure 1.** Changes in tidal flat and eel grass (*Zostera marina*) areas (1945–1996). From the Ministry of Environment, www.env.go.jp and www.biodic.go.jp, accessed July 2019.

To obtain a bird’s-eye view of stocking activity, I first created maps of salmon and marine hatcheries. Approximate locations of salmon hatcheries were plotted on a map of Japan using 15 regional maps of salmon hatchery locations obtained from the website of the Japan Fisheries Research and Education Agency (FRA, salmon.fra.affrc.go.jp). I identified 262 salmon hatcheries in 11 prefectures of northern Japan, among which 242 are privately managed, mainly by fishers and cooperatives (Fig. 2a). The seven prefectural and 13 national hatcheries are primarily research facilities. The major target salmonid species are chum salmon (*Oncorhynchus keta*) and pink salmon (*O. gorbuscha*). The number of released chum salmon in Japan has increased remarkably since the 1970s, reaching a historical maximum of 2.1 billion per annum in 1991. Since then, however, releases of chum salmon have dropped— to 1.5 billion in 2018 (North Pacific Anadromous Fish Commission, NPAFC, 2019) (Fig. A1). A similar increasing trend occurred with pink salmon starting in the 1970s; however, the number of released pink salmon is much smaller, namely, 113 million in 2018, which is approximately 7% of that of chum salmon. The number of masu salmon (*O. masou*) released in 2018 amounted to 7 million. Sockeye salmon (*O. nerka*) are also released, but at even smaller numbers (NPAFC, 2019).

**Figure 2.** Maps showing the locations of marine and salmon hatcheries in Japan operated by different sectors. From FRA (salmon.fra.affrc.go.jp, accessed August 2019) and National Association for Promotion of Productive Seas (http://www.yutakanaumi.jp/, see text).

The locations of marine hatcheries—so-called “saibai-gyogyo centres”—were also plotted by using a list of their addresses compiled by the National Association for Promotion of Productive Seas. Presently, 65 prefectural marine hatcheries operate in 43 prefectures (Fig. 2b). In addition, 12 FRA research stations are operating; these were once national marine hatcheries managed by the Japan Sea-Farming Association, which merged with the FRA following administrative reform in 2003. Because the FRA stopped seed production for release, four previous national marine hatcheries have closed as of 2018. In 2017, 74 marine species (excluding salmon) were being released in Japan, including 34 fishes, 10 crustaceans, 22 shellfishes, and 8 other marine species (e.g., sea urchin, sea cucumber, and octopus). I organized the number of released seeds of the 16 major species whose annual release exceeded million seeds for the period 1983–2017 according to annual seed production and release statistics (Fishery Agency et al. 1985–2019). These species are abalone (*Haliotis* spp.), Japanese scallop (*Mizuhopecten yessoensis*), Manila clam (*Ruditapes philippinarum*), green tiger prawn (*Penaeus semisulcatus*), kuruma prawn (*Marsupenaeus japonicus*), offshore greasyback prawn (*Metapenaeus ensis*), swimming crab (*Portunus trituberculatus*), black sea bream (*Acanthopagrus schlegelii*), flatfish (e.g., *Limanda yokohamae*), Japanese flounder (*Paralichthys olivaceus*), Pacific herring (*Clupea pallasii*), red sea bream (*Pagrus major*), sailfin sandfish (*Arctoscopus japonicus*), tiger puffer (*Takifugu rubripes*), sea urchins (e.g., *Strongylocentrotus intermedius* and *Heliocidaris crassispina*), and sea cucumber (*Apostichopus armata*). The number of releases of 10 species has been decreasing, including iconic target species such as kuruma prawn, swimming crab, abalone, red sea bream, Japanese flounder, and sea urchin (Figs. A2, A3; Supplementary Data). The number of releases of Japanese scallop has slightly increased, while releases of tiger puffer, Pacific herring, and sea cucumber have significantly increased. The changes in release practices reflect changes in subsidies and budgets of the Fisheries Agency and prefectural governments, thus showing that the Japan marine stock enhancement programme has been governmentally led. The only exception is Japanese scallop stocking, which is run by the fishers themselves.

### Evaluation of stocking effects

In earlier surveys, the counting of external tags reported by fishers was the main method used to estimate recapture rates (Kitada 1999). This type of survey was applied to estimate migration, growth, and life history, including comparisons of seed quality (Kitada & Hirano 1987; Shiota & Kitada, 1992; Kitada et al. 1994; Okouchi et al. 1994; Takaba et al. 1995). Tag-reporting rates were often very small, however, and tag-shedding and tagging mortality caused biased estimates. Researchers became aware of bias in the results when they estimated stocking effectiveness. As an alternative approach, surveys of commercial landings at fish markets (SCFMs) have been used to estimate landings of released seeds and the hatchery contribution to the landings. By sampling landings at fish markets, researchers have aimed to comprehensively estimate the stocking effect. The SCFM approach was first applied to red sea bream in Kagoshima Bay. To identify released fish at this location, almost all red sea bream landed at the Kagoshima Fish Market were checked for a deformity of the internostril epidermis (Shishidou 2002; Shishidou & Kitada 2007; Kitada et al. 2019). To sample Japanese flounder in Miyako Bay, all flounder at the Miyako fish market were examined (Okouchi et al. 1999, 2004). Because such a census approach was generally difficult to apply, sampling surveys have been accordingly introduced throughout the country. The procedure used in these surveys is to check all fish landed and sold at a selected market on a selected day to differentiate released seeds from wild individuals. This latter approach, treated as a procedure, implemented in two stages to allow for the formulation of estimators and their variance (Kitada et al. 1992), has been applied to abalone (Kojima 1995), kuruma prawn (Yamaguchi et al. 2006), and masu salmon (Miyakoshi et al. 2001). When information on total landing days is lacking, simple random sampling is assumed, and landings from releases are then estimated (Obata et al. 2008). Some other SCFM-based studies have assumed simple random sampling, but the precision was not evaluated in most cases. In yet another approach, genetic marking has been applied to mud crab (*Scylla paramamosain*) to estimate recapture rates using genetic stock identification (Obata et al. 2006).

A population dynamic model was developed to predict the effects of fishing regulations and hatchery releases on fishery production. Under the auspices of the Fisheries Agency, the Japan Sea-Farming Association tested this model using red sea bream and Japanese flounder data collected throughout the country (Kitada & Okouchi 1994). The model parameters were then adjusted to fit predicted catches with observed ones. Although the retrospective prediction was particularly sensitive to natural mortality coefficient values and parameters in the reproduction models, such models of simulated fishery production are convenient for ascertaining the relative effect of fishery management and release strategies. Indeed, similar simulation models are presently being used for Pacific herring, Japanese flounder, and tiger puffer. In many cases, however, the predictions have failed—such as with Japanese Spanish mackerel (*Scomberomorus niphonius*) in the Seto Inland Sea (Obata et al. 2007) and red sea bream in Kagoshima Bay (Shishidou et al. 2012).

To boost the effectiveness of hatchery releases, a marine-ranching project led by the Fishery Agency endeavoured to shape ‘natural’ farms to provide sound habitat, such as enhanced seaweed communities to promote nutrient enrichment in deep-sea upwelling systems (AFFRC 1989). The major technologies applied for marine fishes that can be reared and released as seeds around artificial reefs have been hatchery releases and artificial reef construction, and feeding systems with acoustic conditioning are also a popular tool (Kudo & Kimoto 1994; Kayano et al. 1998). I could not find scientific literature evaluating the effectiveness of Japan’s marine-ranching projects, and their usefulness thus remains largely unknown. Consequently, I excluded the effects of marine ranching in Japan from my subsequent analyses.

## Meta-analysis of stocking effects

### Indices of stocking effect and Bayesian summary statistics

I used recapture rates, yield-per-release (YPR), and economic efficiency as indices of the performance of hatchery releases. YPR is defined as the weight of fish caught (g) per individual released (Svåsand et al. 2000; Kitada & Kishino 2006; Hamasaki & Kitada 2008b) as follows:

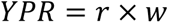

where *r* is the recapture rate of released juveniles, and *w* (g) is mean body weight per recaptured individual. Economic efficiency (*E*) was calculated as the ratio of net income to release costs:

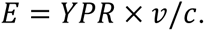

Here, *v* is fish price per gram, and *c* is the cost of each seed; therefore, *v*/*c* is the cost performance of a seed (Kitada 2018). The true mean for the recapture rate, YPR, and economic efficiency for each study (*y_i_*) was not observed but instead estimated (*ŷ_i_*,*i* = 1,…, *n*). When samples (i.e., cases within a study) were taken by random sampling, then E(*ŷ*_*i*_) = *y*_*i*_, and the variance within each study 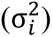 was also estimated from the data 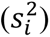.

The release variables, such as tags, sizes at release, release areas, and time periods of surveys, varied in each study; therefore, the summary statistics needed to account for this variation among studies. Here, I assumed a superpopulation for the recapture studies by applying a Bayesian approach. The sample mean of the *i-*th case study (*ŷ*_i_) may follow a normal distribution in accordance with the central limit theorem. Because studies were carried out in different areas and years, each study could be regarded as independent. For this condition, the approximate likelihood of the sample mean *ŷ_i_*(*i* = 1,…, *n*) can be written as:

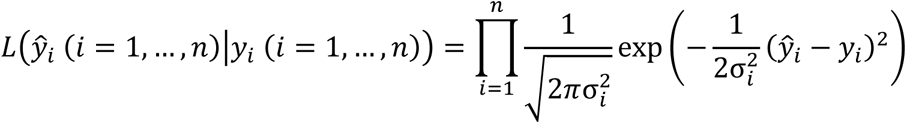

I assumed a prior distribution for *y_i_*(*i* = 1,…, *n*), where *y_i_* is assumed to be normally distributed around the mean *μ* with variance *σ*^2^. The mean and variance of the superpopulation are hyperparameters. The prior distribution of *y*_*i*_ (*i* = 1,…, *n*) can be written as:

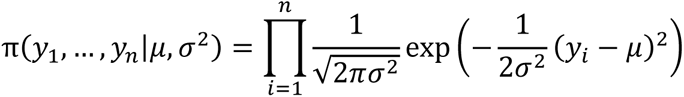

By integrating the product of the prior distribution and the likelihood function in terms of *y_i_*, the marginal likelihood function was derived. In this case, it was explicitly obtained as:

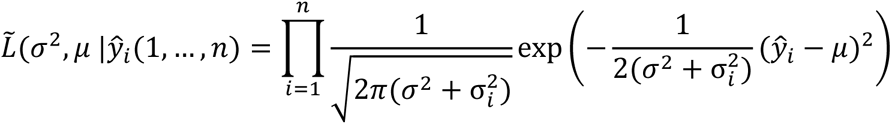

According to this equation, the sample mean *ŷ_i_* is distributed around the mean *μ*, with variances calculated for between cases and within cases.

The log marginal likelihood function is:

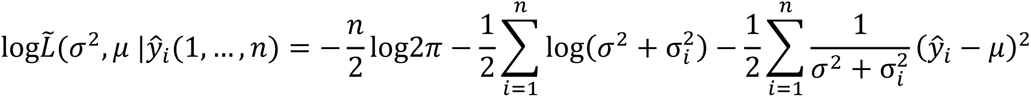

The first derivatives on *μ* and *σ*^2^ are then:

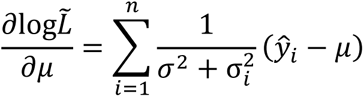

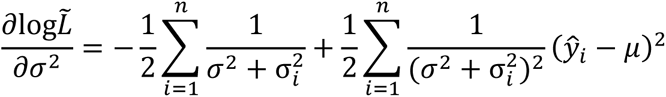

From ∂log*L͂*/∂*μ* = 0, the maximum likelihood estimator (MLE) of *μ* as a weighted average of *ŷ_i_* (*i* = 1,…, *n*) is obtained as follows:

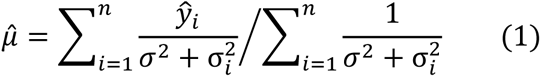

By substituting *σ*^2^ with 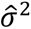 from Eq. (2) and 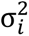 with the sample variance 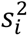, the MLE of *μ* (posterior mean) is obtained.

Assuming for simplicity that 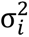 does not depend on a particular case, 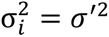 for all *i* (1,…, *n*) and from ∂log*L͂*/∂*σ*^2^ = 0, the MLE of *σ*^2^ is:

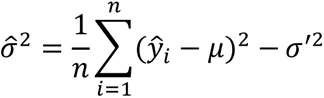

By estimating *μ* by the weighted average 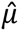 given by Eq. (1) and *σ*^′2^ by the mean of the sample variance 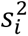, the approximate variance estimator is:

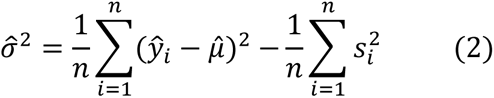

The MLE of *μ* used here had a different weight than the weighted mean (also the MLE) used in my previous study (Kitada 2018), while 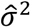 was the same. Eq. (1) provided a more realistic estimate than that obtained using the previous weighted average, particularly when very small *ŷ_i_* values (*ŷ_i_*<1) were included (i.e., when there was large variation in *ŷ_i_*). The average and standard deviation (SD) with a 95% confidence interval (mean ± 2 SD) were visualized for each case using the ‘forestplot’ function in R. When the lower confidence limit took a negative value, it was replaced by 0.

### Recapture rate

I summarized 21 full-scale programmes that reported recapture rates of hatchery individuals based on various marking methods (Table A1). Eighteen of these programmes were also included in my previous study (Kitada 2018). The mean recapture rate of chum salmon was newly calculated for Hokkaido, the main production area, and was based on simple return rates (number of fish returned after 4 years/number released) taken from the FRA website (salmon.fra.affrc.go.jp). Recapture rates from 17 of 21 studies of marine species were estimated using SCFM data or a combination of SCFM data and reported recaptures. In addition, recapture rates were summarized from previous reviews for the whole country for red sea bream (Kitada & Kishino 2006), kuruma prawn (Hamasaki & Kitada 2006, 2008b), and abalone (Hamasaki & Kitada 2008a). Recapture rates were calculated for black rockfish (*Sebastes schlegelii*) (Nakagawa et al. 2004), Japanese flounder (Kitada et al. 1992; Tominaga & Watanabe 1998; Ishino 1999; Atsuchi & Masuda 2004; Tomiyama et al. 2008), Japanese scallop (Kitada & Fujishima 1997), Japanese Spanish mackerel (Obata et al. 2008), mud crab (Obata et al. 2006), short-spined sea urchin (*Strongylocentrotus intermedius*) (Sakai et al. 2004), spotted halibut (*Verasper variegatus*) (Wada et al. 2012), swimming crab (Okamoto 2004), and tiger puffer (Nakajima et al. 2008). Moreover, I newly added four cases of three species: barfin flounder (*Verasper moseri*) in Hokkaido (Koya 2005; Murakami 2012; NPJSEC 2015), red spotted grouper (*Epinephelus akaara*) in Osaka Bay (Tsujimura 2007), and tiger puffer in the Ariake Sea (Matsumura 2005, 2006); these cases were included because stocking effects have recently been reported for the first two species and the number of tiger puffer individuals released has increased in recent years (Fig. A3). In particular, more than one million juveniles (∼8 cm total length, TL) of barfin flounder in Hokkaido have been released annually along the Pacific coast since 2006.

In total, recapture rates were included for 15 species comprising eight marine fishes, one salmonid, three crustaceans, two shellfishes, and one sea urchin (22 programmes) (Table A1). After excluding from analysis any cases that reported only point estimates, the recapture rates of marine species from 20 programmes (Table A1) varied widely among species and cases, ranging from 0.9% to 34.5% (Fig. 3). Japanese scallop had the highest recapture rate (34.5 ± 10.2%), followed by sea urchin (18.2 ± 17.5%), both with relatively large variations. Most of the marine fishes, namely, barfin flounder, black rockfish, Japanese flounder, Japanese Spanish mackerel, red spotted grouper, spotted halibut, and tiger puffer (to age 0+), had recapture rates in the range of 11%–15%. Abalone likewise had a relatively high and highly variable recapture rate (12.2 ± 8.1%). Red sea bream had recapture rates of 7%–8% with only small variation. In contrast, the return rate of chum salmon was much smaller, 3.6 ± 1.1% on Hokkaido and 1.6 ± 0.6% on Honshu (figure not shown). Crustaceans generally had values smaller than 5%, with large variations in recapture rates observed for kuruma prawn (2.8 ± 4.5%) and mud crab (0.9 ± 0.7%). The posterior mean recapture rate was 8.3 ± 4.7%. The empirical distribution of the posterior mean showed that the recapture rates of any species could fall into that range with 95% probability.

**Figure 3.** Forest plot of recapture rates from large-scale hatchery releases in Japan. Thin lines indicate 95% confidence intervals, with arrows (in the case of Japanese scallop and sea urchin) indicating that the confidence intervals penetrate the scale. Areas of the squares are proportional to the weight of the mean.

### YPR and economic efficiency

To analyse YPR and economic efficiency, I revisited 10 cases for which YPR and economic efficiency values were previously reported (Kitada 2018) and added six new cases: those of barfin flounder in Hokkaido, Japanese flounder at Fukushima, red spotted grouper in Osaka Bay, tiger puffer in the Ariake Sea, and swimming crab in Lake Hamana and in the Seto Inland Sea (Table A2). YPR and economic efficiency values were recalculated for kuruma prawn (Hamasaki & Kitada 2006, 2008b), swimming crab (Hamasaki et al. 2011), and abalone (Hamasaki & Kitada 2008a) based on previous reviews. For the calculation of the YPR of chum salmon, mean body weights were revised from 1974–2017 catch statistics obtained from the NPAFC and 1974–2017 return rates in Hokkaido obtained from the FRA (http://salmon.fra.affrc.go.jp/zousyoku/sakemasu.html). The revised mean body weight of chum salmon was 3 197 ± 306 g. Mean body weights were also recalculated for Japanese scallop based on Kurata (1999), yielding a revised estimate of 177 ± 30 g. YPR values reported for pink salmon in Hokkaido by Ohnuki et al. (2015) were given in monetary terms (1.5–2.2 yen/released individual), which was calculated from the estimated proportion of Hokkaido-originated hatchery fish in the landings. Because the YPR of pink salmon in Hokkaido was not provided in terms of weight, recapture rate, or cost performance of a seed (*v*/*c*), I excluded this case from the analysis.

YPR and economic efficiency were ultimately evaluated for 16 cases involving 12 species (Table A2). After omitting cases with only point estimates for YPR and/or those without information needed for calculation of *v* and *c*, a meta-analysis was performed on 12 cases involving 9 species for YPR and 13 cases involving 10 species for economic efficiency. YPR values varied between species and cases. The empirical distribution of the summary statistics had a long tail to the right, with a posterior mean of 65 ± 74 g (Fig. 4a). YPR was highest in barfin flounder (182 ± 9 g) and Japanese Spanish mackerel (170 ± 8 g), followed by chum salmon (119 ± 45 g) and Japanese scallop (61 ± 18 g), both in Hokkaido. Red sea bream and Japanese flounder had YPR values of 30–59 g. Other YPR values were 34 ± 10 g for swimming crab, 26 ± 19 g for abalone, 4 ± 3 g for mud crab, and 0.9 ± 1.5 g for kuruma prawn.

**Figure 4.** Performance of large-scale hatchery releases in Japan. Forest plots of **(a)** yield-per-release (YPR), **(b)** economic efficiency, and **(c)** YPR vs. economic efficiency. Note: the seed cost used in the analysis did not include personnel expenses, facilities, monitoring, or administration costs. SIS = Seto Inland Sea.

The posterior mean of economic efficiency was 2.8 ± 6.1. The economic efficiency of several cases ranged from approximately 1 (the break-even point) to 2, with the lower 95% confidence limit below 0 (Fig. 4b). The highest economic efficiencies were those of chum salmon (19 ± 7) and Japanese scallop (18 ± 5) in Hokkaido, where the seed cost for chum salmon was set at 2.5 yen/juvenile (Hokkaido Salmon Propagation Association 2017) (Table A2). The economic efficiency of red sea bream in Kagoshima Bay was also high (5 ± 3), with similar economic performances noted for abalone across coastal Japan (4 ± 2) and barfin flounder on the Pacific coast of Hokkaido (3 ± 0.1). Japanese flounder had a much smaller economic efficiency, 0.9–1.6. Crustaceans consistently exhibited economic efficiency values around 1; among them, kuruma prawn had the lowest value, 0.7 ± 0.9. Economic efficiency is a function of YPR (recapture rate × body weight) and the cost performance of a seed (*v*/*c*). As shown in Fig. 4c, which is a scatter plot of YPR vs. economic efficiency that depicts the stocking-performance characteristic of each species, chum salmon and Japanese scallop in Hokkaido had the highest economic efficiencies.

I did not analyse net present value (NPV) (Sproul &Tominaga 1992; Moksness & Støle 1997; Moksness et al. 1998; Svåsand et al. 2000) because various data, such as annual costs for harvest, management, and interest rates, were not available for every case. If the economic efficiency estimates obtained here had relied on NPV, they would have been smaller (Kitada 2018); in that case, the values of the estimates would have depended on interest rates and time duration (although interest rates in Japan are currently low, <0.05%). In addition, the seed cost used in the analysis did not include personnel expenses, facilities, monitoring, or administration costs; furthermore, it did not account for the cost of negative effects on natural populations and ecosystems (Waples 1991; Winton & Hilborn 1994; Hilborn 1998; Waples 1999; Waples & Drake 2004; Amoroso et al. 2017). The estimates of economic efficiency obtained here are thus too high. If all the costs were available and included in the analysis, all cases might be unprofitable except Japanese scallop in Hokkaido, where fishers pay most of the programme costs. Chum salmon in Hokkaido had higher economic efficiency than Japanese scallop, but it has been suffered from continuous decline in the population size since around 1996 (see, later discussion).

### Macro-scale contribution of released seeds to commercial landings

I estimated the contribution of hatchery releases to the commercial catch of eight species (Fig. 5). These iconic species of Japan’s marine stock enhancement and sea ranching programmes are intensively released within the range of Japanese waters. Trends in catches were tested using Mann-Kendall trend test with R function MannKendall. After releases of these eight species, catches of Japanese scallop (*τ* = 0.81, *p* = 2.2 × 10^-16^) and Japanese Spanish mackerel increased (*τ* = 0.80, *p* = 9.5 × 10^-7^) (Fig. 5). Catches of Japanese flounder (*τ* = 0.06, *p* = 0.614) were stable and those of red sea bream were slightly increasing (*τ* = 0.33, *p* = 0.005) . In contrast, catches of kuruma prawn (*τ* = −0.95, *p* = 3.9 × 10^-15^) and swimming crab (*τ* = −0.66, *p* = 1.4 × 10^-8^) have decreased continuously since the mid-1980s. Continuous declines in abalone (*τ* = −0.84, *p* = 8.4 × 13) and sea urchin (*τ* = −0.89, *p* = 2.2 × 10^-14^) catches have also been observed since the early 1970s. Different reasons have been advanced to explain these various trends. Cases showing increasing catch levels are consistent with the frequent claims of management agencies and hatchery advocates that the practice of hatchery release should be successful. A popular explanation for cases displaying a stable catch is that hatchery release stabilizes recruitment. Instances of decreasing catches have been attributed to decreased numbers of releases; if more seeds were released, catch levels would increase. To test these ideas, estimates of the contribution of hatchery-released seeds to the catches would be helpful.

**Figure 5.** Total catch and recovery from releases of representative species in Japanese stocking programmes for all of Japan. Vertical lines depict recovery (expected catch from releases), which were estimated by multiplying values of yield-per-release (YPR) (see Table A2) and the numbers released (Supplementary Data). For sea urchin, no YPR data were available.

To evaluate hatchery contributions to major species for all of Japan, I computed the expected catch from released seeds by multiplying the average YPR (listed in Table A2) by the number of fish released every year (Supplementary Data). The contribution to Japanese Spanish mackerel was calculated solely from Seto Inland Sea data because hatchery releases of this species in Japan are made only at that location. This simple analysis assumed that YPR was constant over years and that released seeds created the catch in the same year. The analysis thus did not account for a time lag; however, it allowed macro-scale comparisons between species of the approximate contribution of hatchery releases (Kitada & Kishino 2006).

The largest proportion of released seeds, 76 ± 20%, was for Japanese scallop, with released spat having a stable contribution (Fig. 5, vertical bars; Fig. A4). Wild scallop created by natural reproduction also contributed to the total catch. The catch decreased substantially during 2015–2017 following a bottom disturbance off the Okhotsk coast caused by a low-pressure bomb in December 2014 (Kitada 2018); according to the estimates for that time period, almost all of the catch comprised released spat, thus indicating that scallop populations in the fishing grounds were heavily damaged by the bottom disturbance. Although the analysis was simple, the results demonstrate that this approach was able to describe the population dynamics of Japanese scallop in Hokkaido.

The hatchery contribution to the increased catch of Japanese Spanish mackerel was very small, 2 ± 2%. The catch of this fish continued to recover after releases; however, the number of released juveniles was reduced 10 years after the beginning of the release project. These results clearly indicate that the population dynamics of this species were unaffected by the releases. A previous study found 35% variation in the biomass of age-0 Japanese Spanish mackerel, an observation that could be explained by the biomass of a prey fish, Japanese anchovy (Nakajima et al. 2013). In another investigation, genetic stock identification following releases in 2001 and 2002 found admixture proportions of hatchery-origin fish at 8%–15% (Nakajima et al. 2014). Interestingly, the genetic admixture contribution of hatchery fish in the Seto Inland Sea was much higher than estimated contribution rates (2 ± 2%) in the present study, similar to findings for red sea bream in Kagoshima Bay (Kitada et al. 2019), again implying a trans-generational genetic effect. The hatchery contribution of Japanese flounder was 11 ± 4%, and that of red sea bream was 7 ± 2%. Over two decades, the number of released juveniles decreased by ∼50% for flounder and ∼63% for red sea bream, but catches of wild fish remained stable and/or slightly increased; this indicates that the hatchery releases did not boost the population size of either species on a macro-scale. On the basis of carrying capacity, natural reproduction should have supported the recruitment.

Among crustaceans, the hatchery contribution of kuruma prawn was 13 ± 5%. These results are in agreement with estimates for kuruma prawn from previous research (Hamasaki & Kitada 2013), namely, ∼10% throughout Japan from 1977 to 2008. In that study, the decline in kuruma prawn catches was potentially attributed to warming ocean conditions, decreased fishing efforts due to fewer fishers, and reduced hatchery releases. A continuously decreasing trend in the catch implies a decline in natural recruitments, which suggests an environmental effect is responsible (Fig. 5). Reduced fishing efforts would likely work to increase population size, as demonstrated in the case of red sea bream (Kitada et al. 2019). A reduction in fishing efforts can therefore be excluded as a cause of the decreasing catches of kuruma prawn. For swimming crab, the hatchery contribution was 20 ± 5%; the catch continuously decreased, again showing a decline in natural recruitment. Juvenile crabs are also found on tidal flats after the C4 stage (∼16 mm carapace width, CW) and/or C5 stage (∼22 mm CW) (Hamasaki et al. 2011). The catches of swimming crab were positively correlated with those of kuruma prawn (*r* = 0.52, *t* = 4.49, df = 53, *p* = 3.9 × 10^-5^), suggesting a common environmental effect on juveniles of both species.

Abalone displayed a relatively high and stable hatchery contribution rate (28 ± 10%), yet the catch of this species continued to decrease, indicating decreases in natural recruitment. A negative correlation has been found between the Aleutian Low-Pressure Index, winter sea surface temperatures, and catches of Ezo abalone (*Haliotis discus hannai*) in northern Japan (Nakamura et al. 2005; Hayakawa et al. 2007), and cold winter seawater temperatures (<5 °C) affect the survival of young Ezo abalone (Takami et al. 2008). Seaweed community richness (carrying capacity) is the key factor for successful abalone stocking in Japan (Hamasaki 2008). I could not calculate the hatchery contribution rates of sea urchin because no published information for calculating YPR values was found. Surprisingly similar to abalone, however, the catch of sea urchin exhibited a decreasing trend (*r* = 0.96, *t* = 24.63, df = 53, *p* = 2.2 × 10^-16^), thereby indicating that sea urchin abundance likewise heavily depends on the richness of the seaweed community.

### Economic, ecological and genetic impacts of the world’s largest programmes

Broodstock management for hatchery stocking typically involves three different approaches: (I) wild collection of larvae, (II) wild collection of parents (may include individuals of hatchery-origin), and (III) captive breeding (Kitada 2018). Among the 22 cases examined above, 15 were type III, 6 were type II, and 1 was type I (Table A1). To summarize the impacts of the world’s largest stocking programmes, I analysed three programmes with contrasting management approaches and with the highest economic efficiency values: red sea bream in Kagoshima Bay and Japanese scallop and chum salmon, both in Hokkaido. The broodstock management approaches used in these programmes are as follows: for Japanese scallop, collection of wild larvae from the sea (type I); for chum salmon, collection of parents from the wild (type II); and for red sea bream, captive breeding in a concrete tank (type III). Because all three broodstock management approaches are thus represented, the three case studies analysed here may be useful for predicting the long-term impacts of hatchery stocking programmes worldwide.

### Red sea bream in Kagoshima Bay (type III)

The hatchery-release programme for red sea bream in Kagoshima Bay is one of the world’s largest programmes for marine fish species and the best-monitored one in Japan. Since the programme’s beginning in 1974, ∼27 million hatchery-reared red sea bream (6–7 cm TL) have been released in the bay. Starting in 1974, the broodstock of red sea bream intended for release in Kagoshima Bay have been reared in a 100-m^3^ concrete tank. In particular, approximately 130 non-local red sea bream broodstock are maintained in the concrete tank for natural spawning and have been repeatedly used as broodstock. In 1999 and 2014, 395 wild, 33 farmed, and numerous 2-year-old hatchery fish produced from the broodstock were added to the broodstock. The number of parent fish used for seed production has varied from approximately 45 to 187 annually. During 1989–2015, 1.6 million red sea bream were examined at fish markets to identify hatchery fish caught in the bay. The catch of hatchery fish reached 126 tonnes by 1991 but thereafter consistently decreased, dropping to 3 tonnes by 2016. This decrease was due to a decline in fitness of the hatchery-reared fish in the wild, which was caused by the repeated use of parent fish reared in captivity (captive breeding) since 1974 (∼nine generations). In contrast, the catch of wild fish increased after 1991 and reached a maximum in 2016 following releases amounting to 168 tonnes. Denser seaweed communities and reduced fishing efforts were the primary factors leading to the recovery of the wild population of red sea bream. These results clearly show that the recovery of nursery habitats and reductions in fishing efforts were more effective than hatchery stocking for recovering depleted populations (see Kitada et al. 2019 for detailed information).

As seen in this example, the hatchery releases of red sea bream into Kagoshima Bay substantially increased fisheries production for the first 15 years, a period during which the programme came to be regarded as representative and successful in Japan; importantly, however, the catch of hatchery fish then steadily decreased and remained very low. The declining catch of red sea bream in Kagoshima Bay was attributed to genetic effects (Kitada et al. 2019) through unintended domestication/selection by captive breeding (Ford 2002; Araki et al. 2007, 2008). Ample evidence exists that captive breeding of salmonids can reduce the fitness of hatchery salmon in the wild (Reisenbichler & McIntyre 1977; Fleming et al. 2000; McGinnity et al. 2003; Araki et al. 2007; Christie et al. 2014). Cumulative recapture rates of red sea bream until age 8+ decreased 15.6 ± 1.7% (standard error, SE) per year. In addition, the rate of fitness reduction in hatchery-reared populations was cohort-specific; it was constant over time within the cohort but exponentially decreased with the duration of captivity (Kitada et al. 2019). Indeed, the proportion of hatchery fish in landings in inner and central parts of Kagoshima Bay and outside of the bay were highest in 1990, at 83.3%, 33.5%, and 7.4%, respectively, but the proportion in 2015 was only ∼1.0% (even in the inner part) (Kitada et al. 2019). This observation suggests that once a fitness decline arises in a hatchery population, the reduction in fitness may continue until the broodstock are completely replaced by wild fish.

Gene flow between red sea bream populations was very high, thereby expanding the area affected by the hatcheries. Although more areas would thus be affected, the influence on the meta-population was not very strong. A genetic diversity analysis of fish collected from Kagoshima Bay suggested that the genetic effects of hatchery releases were gradually diluted by backcrossing with wild populations (Kitada et al. 2019), which would diminish the genetic effects of captive breeding if such effects were additive (Roberge et al. 2008). The increasing population of wild fish in Kagoshima Bay showed no fitness decline attributable to the small proportion of hatchery fish in the meta-population. The genetic effects of captive breeding can be gradually diluted if the proportion of hatchery fish in the recipient population is not substantial and/or hatchery releases stop.

Care should be taken, however, when speculating about cases with very high proportions of hatchery fish in the recipient population, particularly cases that use captive breeding. Captive-reared parents have been repeatedly used for abalone, barfin flounder, black sea bream, Japanese flounder, and tiger puffer (Table A1). In the case of barfin flounder, one million juveniles (∼8 cm TL) were released annually on the Pacific coast of Hokkaido. The catch increased markedly, from 0.01 tonnes in 1996 to over 100 tonnes in 2015, and almost all catches in the North Pacific, from Ibaraki Prefecture to Hokkaido (170 tonnes), consisted of hatchery fish (NPJSEC 2015). In the case of barfin flounder, ∼500 parent fish have been used for seed production for 1–3 generations in captivity, and the expected heterozygosity is very high, 0.87 (Andoh et al. 2013). More generations of barfin flounder are needed before the impacts of this case can be determined.

### Japanese scallop (type I)

Japanese scallop accounts for ∼27% of landings at the Hokkaido fishery (www.pref.hokkaido.lg.jp), where all scallop landings are obtained from releases of wild-born spat or from naturally-reproduced spat of released individuals (Kitada et al. 2001; Nishihama 2001; Uki 2006). Initiated by fishers, who pay most of the programme costs (Uki 2006), the scallop-ranching programmes in Hokkaido are managed by cooperatives. The management system for Japanese scallop used by the cooperatives has four components: (1) mass-releases of wild-born spat (wild-born sea-collected larvae are reared in net cages for 1 year prior to release); (2) removal of predators, such as starfish, before release; (3) monitoring of the density of scallops in the fishing ground; and (4) rotation of fishing grounds chosen for harvesting (Goshima & Fujiwara 1994; Kitada et al. 2001; Uki 2006). The fishing grounds are generally partitioned into four areas, and 1-year-old wild-born spat (∼4.5 cm) reared in net cages are released into a given area after removal of starfish. After 3 years, the 4-year-old released scallops are harvested. The following year, spats are released into another area. This fishing-ground rotation system enables complete prohibition of the fishery for 3 years after release.

Catches of Japanese scallop in the release areas, which have increased remarkably since the first releases in the 1970s, reached a historical maximum of 359 000 tonnes in 2014 (Fig. 6a, updated from Kitada 2018). In 2015, the catch dropped substantially, to 232 000 tonnes (65%), because of a bottom disturbance on the Okhotsk coast caused by a low-pressure bomb on 16–17 December 2014 (Kitada 2018), but recovered to 305 000 tonnes in 2018, as expected. The catch recovery in 2018 would have been created by spat released in 2015. The long-term trends in these release and catch statistics demonstrate that sea ranching of scallop in Hokkaido has been successful, with the highest observed economic efficiency (18 ± 5).

**Figure 6.** Long-term release and catch (return) statistics for **(a)** Japanese scallop (updated from Kitada 2018) and **(b)** chum salmon in Hokkaido (updated from Miyakoshi et al. 2013) (see Supplementary Data).

### Chum salmon (type II)

Chum salmon hatchery-stock enhancement in Japan is one of the world’s largest salmon stocking programmes (Amoroso et al. 2017). Hokkaido produces ∼80% of chum salmon returning to Japan, and fishery production accounts for 21% of the catch at the Hokkaido fishery (www.pref.hokkaido.lg.jp). Because most landings are created by releases of hatchery-born fish (Kaeriyama 1999), fishers pay ∼7% of their landings of chum salmon to the hatcheries in Hokkaido (Kitada, 2014). Long-term trends in releases and catches indicate that the maximum carrying capacity of natural rivers is ∼10 million fish and that a large increase in the fish population was created by hatchery stocks (Fig. 6b, updated from Miyakoshi et al. 2013). Total production in Hokkaido reached a historical maximum of 61 million returns in 2004 and then substantially decreased to 16 million in 2017. Chum salmon in Japan are at the southern margin of the species range. My previous study found that 30% of the variation in decreasing catches could be attributed to an increasing sea surface temperature (SST) anomaly, and 62% was explained by SST and catches by Russians after 1996 (Kitada 2018). Clear differences have been found, however, between the run timing of chum salmon populations in Russia (Jun–Aug) and Japan (Sep–Nov), and migration routes are also different (Kondo et al. 1965; Morita 2016). These results suggest that the negative correlation between Japanese and Russian catches reported in the previous study was spurious.

No evidence of fitness decline has been reported among hatchery-born and wild-born chum salmon in Japan. The observed genetic effect is instead an altered population structure, with some populations nested across seven and/or eight regional groups (Beacham et al. 2008; Kitada 2014; Sato et al. 2014; Kitada 2018). The run-timing distribution of Hokkaido chum salmon has been altered by the preferential enhancement of early-running fish (returning in September/October), and the late-running population has almost disappeared (Miyakoshi et al. 2013). A lower reproductive success has been observed for early-spawning sockeye salmon in Washington State, USA, and early-emerging juveniles have had relatively low survival in recent years. These observations suggest that the skewed distribution of early spawning in Japan could reduce population fitness during a warming climate (Tillotson et al. 2019).

Almost all chum salmon returning to Japan are hatchery-reared fish (Kaeriyama 1999). These hatchery-reared fish are produced every year from returning adults, and the number of maintained parent fish has been very large (i.e., 15 000–85 000) (Kitada 2018). Even so, naturally spawning chum salmon have been detected in 31%–37% of 238 non-enhanced rivers surveyed in Hokkaido (Miyakoshi et al. 2012) and in 94% of 47 enhanced rivers and 75% of 47 non-enhanced rivers on the northern coast of Honshu Island (Sea of Japan) (Iida et al. 2018). In addition, a study using otolith thermal-marking estimated the proportion of naturally spawned chum salmon to total production at 16%–28% in eight rivers in Hokkaido, with large variation (0%–50%) (Morita et al. 2013). Despite variations in the proportion of natural spawning in rivers, gene flow facilitates the genetic admixture of hatchery-released fish, hatchery descendants, and wild fish in the entire Japan population. To visualize the magnitude of gene flow between populations, I re-examined microsatellite data from 26 chum salmon populations in Japan (14 loci, *n* = 6,028; Beacham et al. 2008) and computed *F*_ST_ values between population pairs based on the bias-corrected *G*_ST_ (Nei & Chesser 1983) using the GstNC function in the R package FinePop1.5.1. This *G*_ST_ estimator provides an unbiased estimate of *F*_ST_ when the number of loci becomes large (Kitada et al. 2017). I then superimposed a diagram in which population pairs with pairwise *F*_ST_ values < 0.01 were connected by lines onto a map of hatchery locations (Fig. 7). The mean pairwise *F*_ST_ was very small (0.007 ± 0.003). The *F*_ST_ threshold of 0.01 was based on the relationship 4*N*_e_*m* = 99, where *N*_e_ is effective population size and *m* is migration rate, corresponding to 99 effective parents migrating between each pair of populations per generation (see Waples & Gaggiotti 2006). Figure 7 depicts substantial gene flow between rivers. The causal mechanisms of population structuring are migration and genetic drift, and differentiation depends on the number of migrants (*N*_e_*m*) (Waples & Gaggiotti 2006; Hauser & Calvalho 2008). Even in Atlantic cod (*Gadus morhua*), a species with high gene flow, temporally stable but significantly differentiated structure can be detected among populations (Hauser & Calvalho 2008). Constant gene flow among populations can create a stable genetic mixture in a meta-population, such as that observed in Pacific herring populations in northern Japan (Kitada et al. 2017). These results suggest that the nested population structure observed across the seven and/or eight regional groups was caused by past translocations (Beacham et al. 2008; Kaeriyama & Qin 2014).

**Figure 7.** Gene flow between Japanese populations of chum salmon. Populations with pairwise *F*_ST_ < 0.01, as estimated from microsatellite data of 26 populations (14 loci, *n* = 6 028; Beacham et al. 2008), are connected by lines.

Hatchery practices might increase the likelihood of chum salmon to stray (Quinn, 1993). I summarized the results of marking studies of chum salmon juveniles in Hokkaido, where 2 028 thousand hatchery-reared salmon juveniles (3.5–4 cm BL, 0.45–5 g) were fin-clipped and/or operculum-clipped and released without rearing between 1951 and 1955 (Sakano 1960). According to the results, 50 ± 22% of 2 085 recoveries were recaptured in the river of their release, and 84 ± 12% were found when recoveries along nearby coasts were included (Supporting Data); hence, the spawning fidelity of hatchery-reared chum salmon was moderate. Estimates of straying can vary largely between specific hatchery releases and rivers, but the genetic integrity of a population can be altered by straying regardless of the strength of the native population’s spawning fidelity (Quinn 1993). Early theoretical work predicted that >99% (50%) of wild genes with additive effects are replaced by hatchery genes in 12 generations (2 generations for 50% of wild genes) in the case of equal fitness between hatchery and wild fish at a stocking rate of 0.5 (Matsuishi et al. 1995; see also Fig. 5 in Kitada et al. 2019). Taking all of these results in consideration, I speculate that most chum salmon returning to Japan are hatchery-released fish or wild-born hatchery descendants. This situation is similar to that of hatchery-reared red sea bream in Kagoshima Bay. The significant decline in the number of returns of Japanese chum salmon may therefore be caused by a fitness decline in populations induced by long-term hatchery stocking.

## Conclusions

All cases of Japanese hatchery releases, except Japanese scallop, are probably economically unprofitable if the costs of personnel expenses, facility construction, monitoring, and negative impacts on wild populations are taken into account. Stocking effects are generally small, while the population dynamics are unaffected by releases but instead essentially depend on the carrying capacity of the nursery habitat. Hatchery rearing can reduce the fitness of hatchery fish in the wild, and long-term hatchery stocking can replace wild genes and cause fitness decline in the recipient population when the proportion of hatchery fish is very high. Short-term uses of hatchery stocking can be helpful, particularly for conservation purposes, but long-term programmes may harm the sustainability of populations. Recovery of nursery habitats and reduction in fishing efforts often outperform hatcheries in the long run.

## Acknowledgements

I thank Hirohisa Kishino, Katsuyuki Hamasaki, Yasuyuki Miyakoshi, Mitsuhiro Nagata, and Shigeki Dan for useful comments on an earlier version of this manuscript. I expression my appreciation to Toshio Okazaki for valuable suggestions, Tomonari Fujita for providing addresses of marine hatcheries, and Reiichiro Nakamichi for help with map construction. This study was supported by the Japan Society for the Promotion of Science Grant-in-Aid for Scientific Research (KAKENHI) No. 18K0578116.

## Data Availability Statement

All source data used are in the public sector, and links to their online sources are specified in the text or in Supplementary Data.

## Supporting Information

Additional Supporting Information may be found in the online version of this article:

## Supporting Data

Catch and release data for major species in Japanese marine stock enhancement and sea ranching programmes and the results of marking experiments with chum salmon conducted in Hokkaido in the 1950s.

## Appendices

**Table A1.**
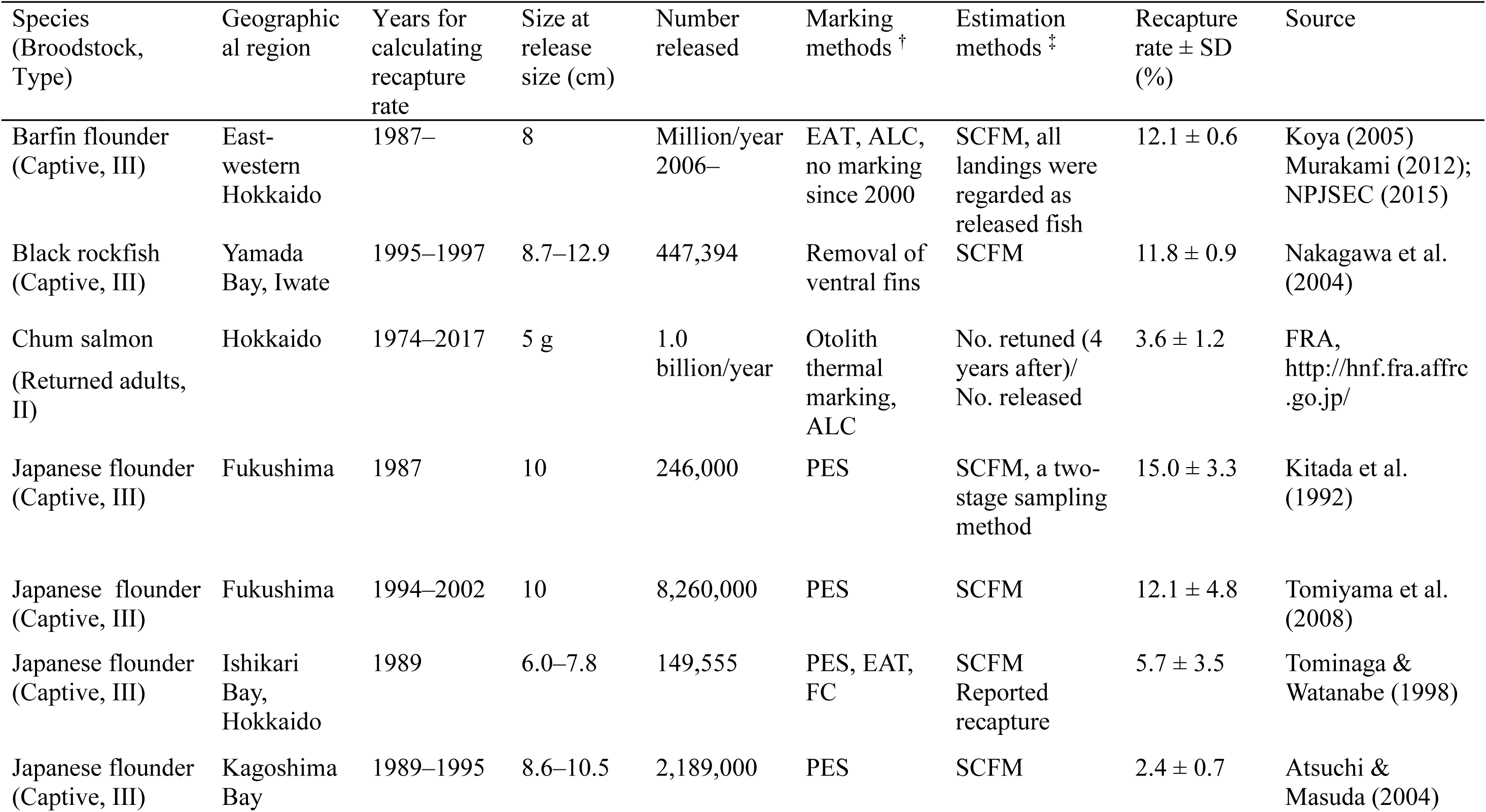
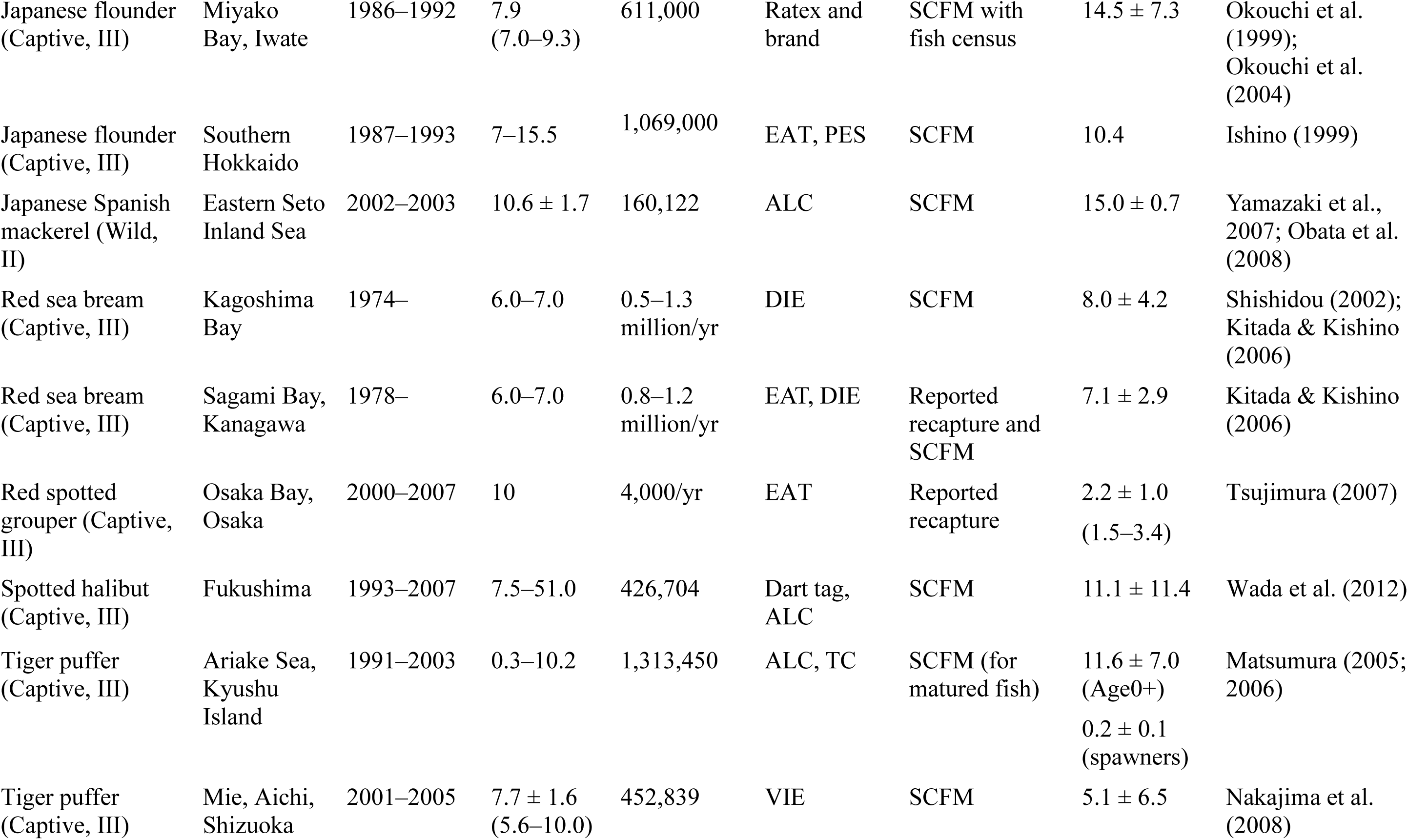
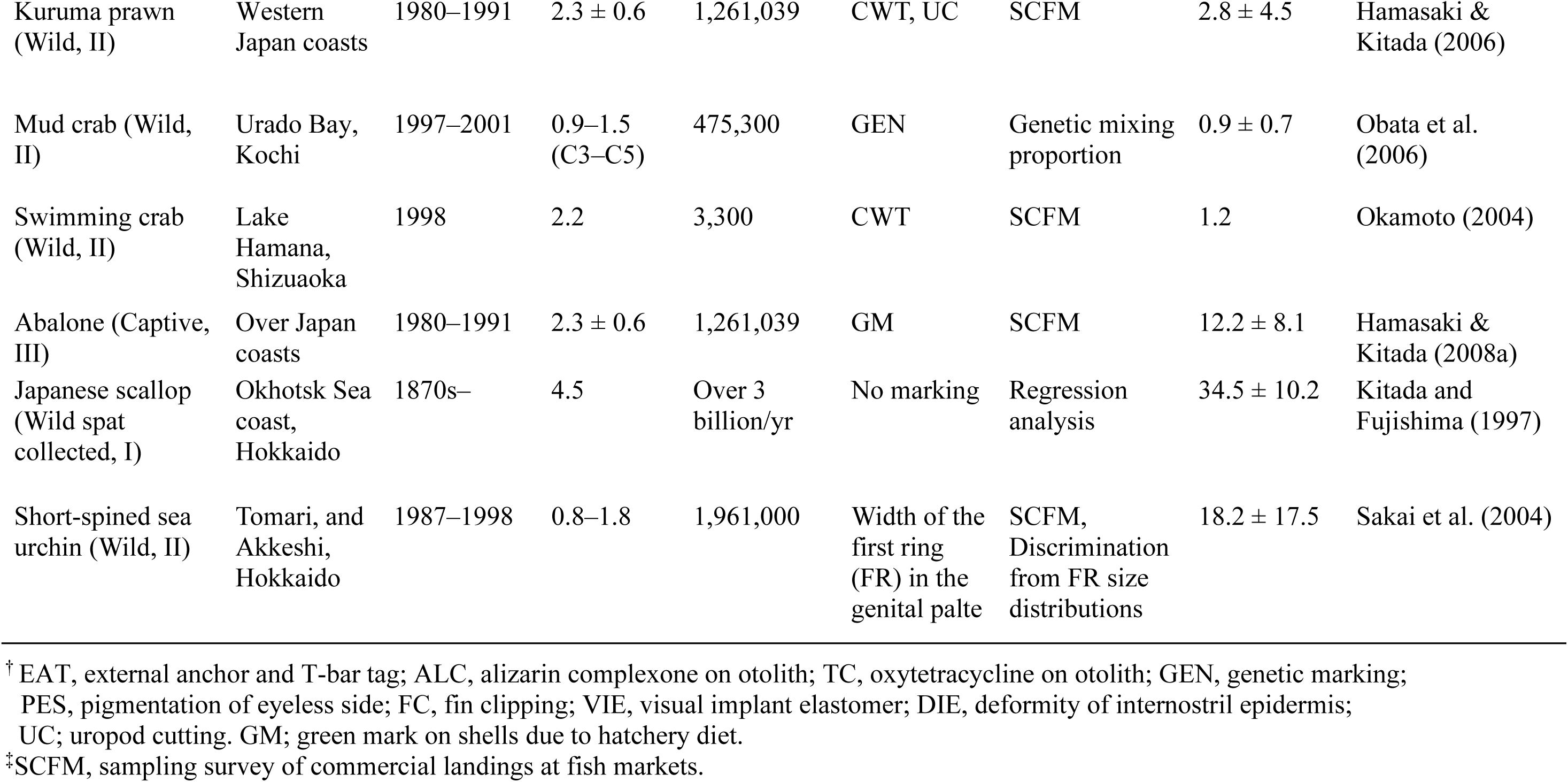
Recapture rates of 21 major marine stock enhancement and sea ranching programmes in Japan

**Table A2.** Performance of 15 major marine stock enhancement and sea ranching programmes in Japan

| Species | Geographical region | Marking methods <sup>†</sup> | Estimation Method <sup>‡</sup> | Recapture rate (%) | YPR <sup>§</sup> (g) | Economic efficiency <sup>¶</sup> | Source |
| --- | --- | --- | --- | --- | --- | --- | --- |
| Barfin flounder | East-western Hokkaido | - | YPR | 12.1 ± 0.6 | 181.5 ± 9.0 | 2.7 ± 0.1 | This study ($w=1500$, $v=1.2$, $c=81$) |
| Chum salmon | Hokkaido | - | YPR | 3.6 ± 1.2 | 118.5 ± 45.1 | 18.9 ± 7.2 | Kitada (2018); This study ($w=3310$, $v=0.4$, $c=2.5$) |
| Japanese flounder | Kagoshima Bay | PES | SCFM | 2.4 ± 0.7 | 29.7 ± 2.6 | 1.1 ± 0.1 | Atsuchi & Masuda (2004); Kitada & Kishino (2006) |
| Japanese flounder | Miyako Bay | PES | CS | 13.5 ± 6.4 | 51.8 ± 24.2 | 1.6 ± 0.7 | Okouchi et al. (2004); Kitada & Kishino (2006) |
| Japanese flounder | Fukushima coast | PES | SCFM | 12.1 ± 4.8 | n.a | 0.9 ± 0.4 | Tomiya et al. (2008) |
| Japanese Spanish mackerel | Eastern Seto Inland Sea | ALC | SCFM | 15.0 ± 0.7 | 169.7 ± 8.3 | 1.0 ± 0.1 | Obata et al. (2008) |
| Red sea bream | Kagoshima Bay | DIE | SCFM | 8.0 ± 4.2 | 59.0 ± 27.2 | 5.0 ± 2.7 | Shishidou (2002); Kitada & Kishino (2006) |
| Red sea bream | Sagami Bay | DIE | SCFM | 7.1 ± 2.9 | 54.9 ± 30.4 | 1.4 ± 0.3 | Kitada & Kishino (2006) |
| Red spotted grouper | Osaka Bay, Osaka | EAT | MR | 2.2 ± 1.0 | 7.7 ± 0.4 | 0.3 ± 0.01 | Tsujimura (2007); This study ($w=350$, $v=5$, $c=150$) |
| Tiger puffer | Ariake Sea,<br>Kyushu Island | ALC, TC | YPR | 11.6 ± 7.0<br>(Age0+)<br>0.2 ± 0.1<br>(spawners) | n.a | 2.4 (Age0+<br>spawners) | Matsumura (2005; 2006) |
| Kuruma prawn | Western coasts | UC,<br>CWT | SCFM | 2.8 ± 4.5 | 0.9 ± 1.5 | 0.7 ± 0.9 | Hamasaki & Kitada (2006,<br>2008b) |
| Mud crab | Urado Bay,<br>Kochi | GEN<br>(mtDNA) | SCFM | 0.9 ± 0.7 | 3.7 ± 3.0 | 1.9 ± 1.5 | Obata et al. (2006),<br>Hamasaki et al. (2011) |
| Swimming crab | Lake Hamana,<br>Shizuoka | CWT | SCFM | 1.2 | 1.5 | n.a | Okamoto (2004) |
| Swimming crab | Seto Inland<br>Sea | - | REG | 17 | 33.6 ± 9.5 | 1.0 ± 0.3 | Hamasaki et al. (2011)<br>This study ($v=0.15$, $c=5$) |
| Abalone | Over coast of<br>Japan | GM | SCFM | 12.2 ± 8.1 | 25.6 ± 19.1 | 3.5 ± 2.4 | Hamasaki & Kitada (2008a) |
| Japanese scallop | Okhotsk coast,<br>Hokkaido | - | YPR | 34.5 ± 10.2 | 60.9 ± 18.0 | 17.9 ± 5.3 | Kitada & Fujishima (1997);<br>Kurata (1999); This study<br>($w=176.5$, $v=0.9$, $c=3$) |

<sup>†</sup>EAT, external anchor and T-bar tag; ALC, alizarin complexone on otolith; TC, oxytetracycline on otolith; GEN, genetic marking; PES, pigmentation of eyeless side; DIE, deformity of internostril epidermis; UC, uropod clipping; CWT, coded-wire tags; GM, green mark on the shell.
<sup>‡</sup>SCFM, sampling survey of commercial landings at fish markets; CS, census of commercial landings; MR, mark-recapture; REG, regression analysis; YPR, yield per release.
<sup>§</sup> Grams of fish caught per individual released.
<sup>¶</sup> Ratio of net income to release cost, excluding personnel expenses, expenditure for hatchery facilities, and monitoring costs;
n.a, not analysed.

**Figure A1.**
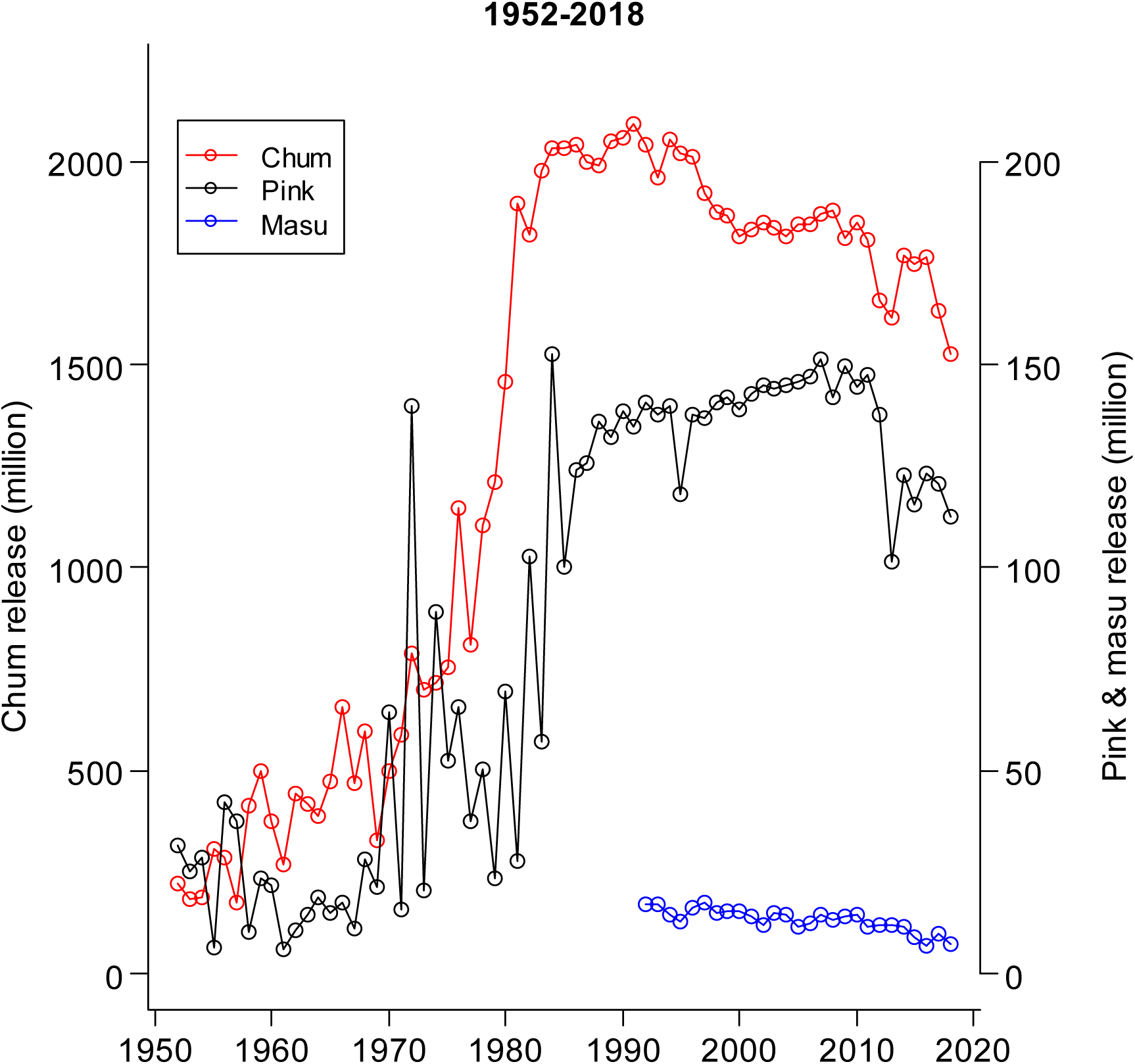
Number of released juveniles of chum, pink, and masu salmon in Japan (1952–2018). Data from the North Pacific Anadromous Fish Commission (www.npafc.org, accessed August 2019).

**Figure A2.**
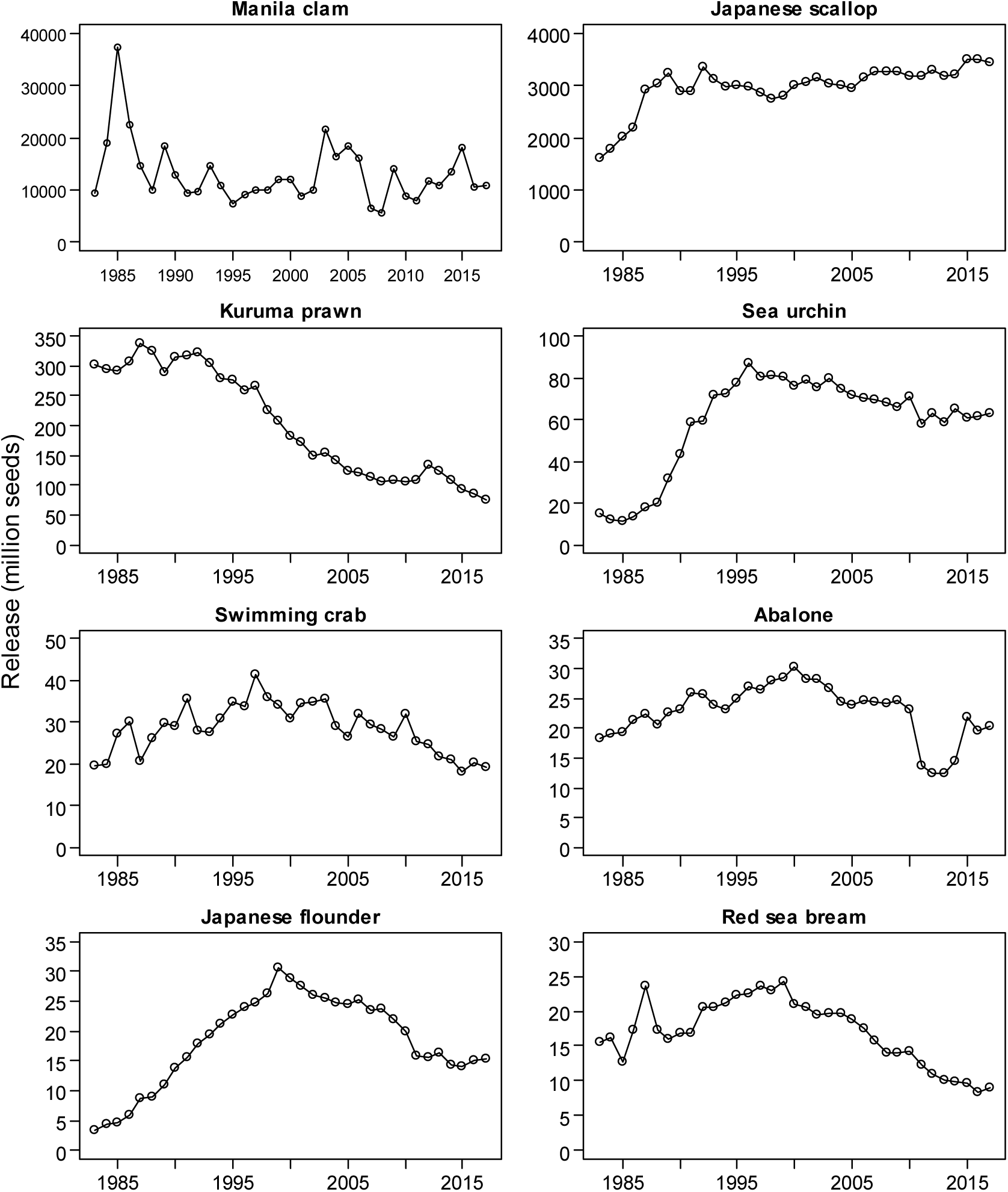
Number of released seeds of the top eight species of Japanese marine stock enhancement and sea ranching programmes (1983–2017). Data from the Fisheries Agency, Fisheries Research and Education Agency, and National Association for Promotion of Productive Seas (1985–2019).

**Figure A3.**
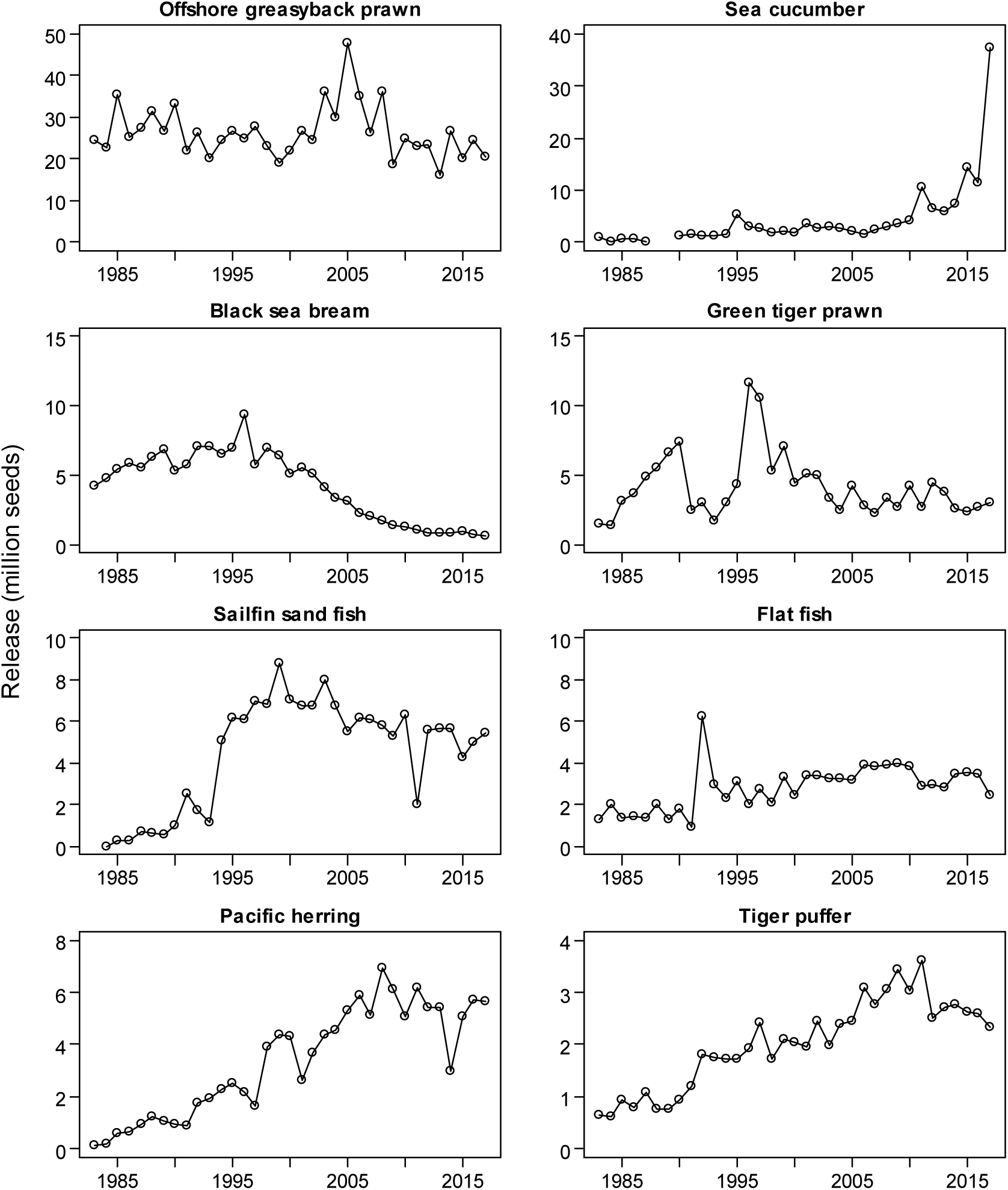
Number of released seeds of 9th–16th-ranked target species of Japanese marine stock enhancement and sea ranching programmes (1983–2017). Data from the Fisheries Agency, FRA, and NAPPS (1985–2019).

**Figure A4.**
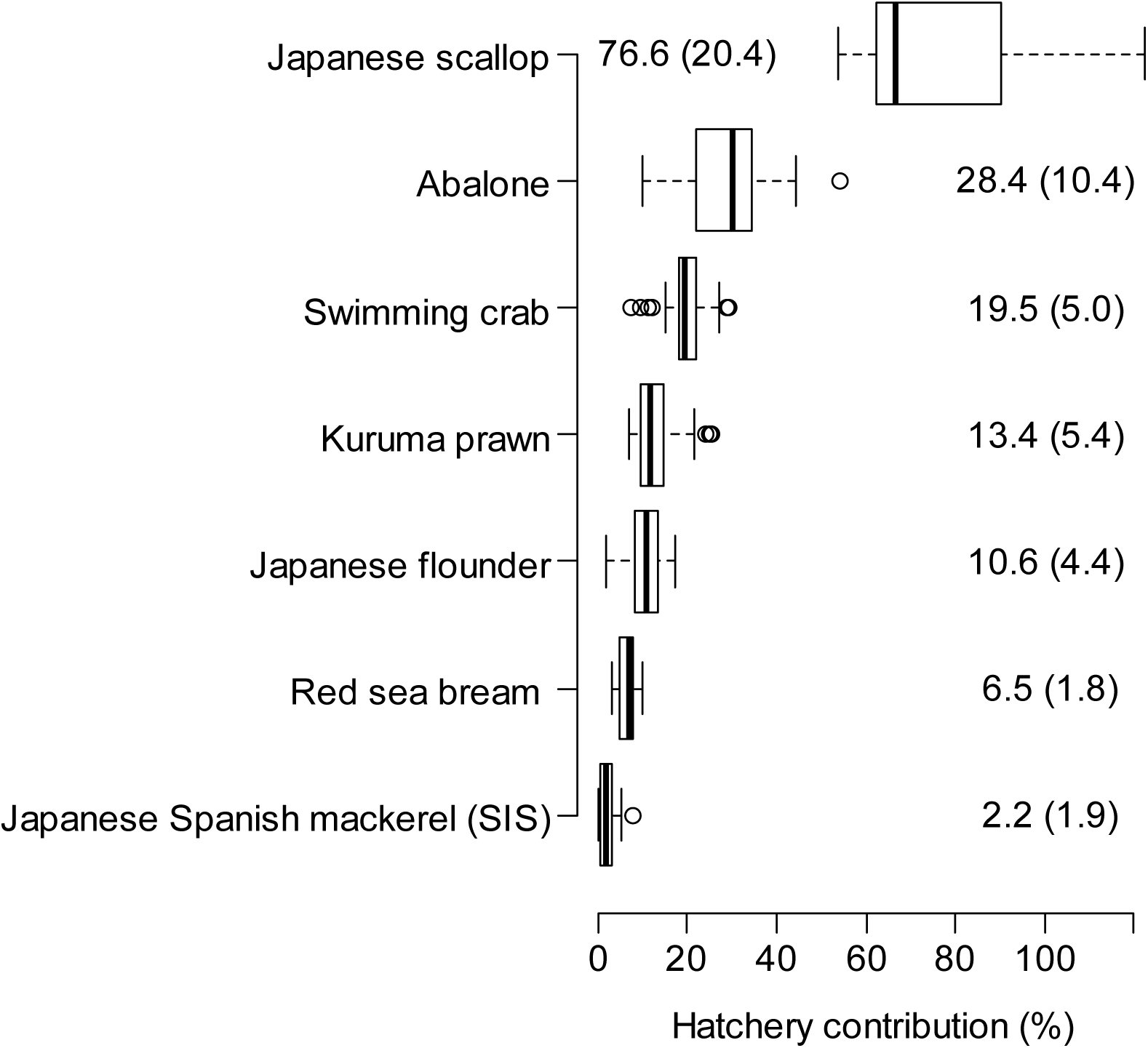
Percent contribution of hatchery-reared individuals to the commercial catch, calculated from Figure 5.

## Notes

#### Summary of Updates

Main text slightly revised with revised title.

## References

1. Agriculture, Forestry and Fisheries Research Council (AFFRC) (1989) Marine Ranching. Koseisha Koseikaku, Tokyo [in Japanese].

2. Amoroso RO, Tillotson MD, Hilborn R (2017) Measuring the net biological impact of fisheries enhancement: Pink salmon hatcheries can increase yield, but with apparent costs to wild populations. Canadian Journal of Fisheries and Aquatic Sciences 74: 1233–1242.

3. Andoh T, Mikasa T, Ichikawa T (2013) Genetic monitoring and conservation programs to maintain the genetic diversity. In: Aritaki M. (ed), Stock Enhancement and Risk Management for Coastal Fishery Resources-Monitoring on genetic diversity and efficiency of stock enhancement, pp. 38–53. Koseisha Koseikaku, Tokyo [in Japanese].

4. Araki H, Cooper B, Blouin MS (2007) Genetic effects of captive breeding cause a rapid, cumulative fitness decline in the wild. Science 318: 100–103.

5. Araki H, Berejikian BA, Ford MJ, Blouin MS (2008) Fitness of hatchery-reared salmonids in the wild. Evolutionary Applications 1: 342–355.

6. Araki H, Schmid C (2010) Is hatchery stocking a help or harm? Evidence, limitations and future directions in ecological and genetic surveys. Aquaculture 308 (Sl): S2–S11.

7. Atsuchi S, Masuda Y (2004) Effectiveness of the releases of hatchery-produced stock of Japanese flounder *Paralichthys olivaceus* in Kagoshima Bay, southern Japan. Nippon Suisan Gakkaishi 70: 910–921 [In Japanese with English abstract].

8. Beacham TD, Sato S, Urawa S, Lei KD, Wetklo M (2008) Population structure and stock identification of chum salmon *Oncorhynchus keta* from Japan determined by microsatellite DNA variation. Fishery Science 74: 983–994.

9. Bell JD, Rothlisberg PC, Munro JL, Loneragan NR, Nash W J, Ward RD et al. (2005) *Restocking and stock enhancement of marine invertebrate fisheries*. Advances in Marine Biology 49, Elsevier, Oxford.

10. Bell JD, Leber KM, Blankenship HL, Loneragan NR, Masuda R (2008) A new era for restocking, stock enhancement and sea ranching of coastal fisheries resources. Reviews in Fisheries Science 16: 1–9.

11. Blaxter JHS (2000) The enhancement of marine fish stocks. Advances in Marine Biology 38: 1–54.

12. Christie MR, Marine ML, French RA, Blouin MS (2012) Genetic adaptation to captivity can occur in a single generation. Proceedings of the National Academy of Sciences 109: 238–242.

13. Christie MR, Ford MJ, Blouin MS (2014) On the reproductive success of early-generation hatchery fish in the wild. Evolutionary Applications 7: 883–896.

14. Fisheries Agency, Fisheries Research Agency (FRA), National Association for the Promotion of Productive Sea (eds.) (1985–2019) Annual Statistics of Seed Production and Release in 1983–2017. FRA, Yokohama [in Japanese].

15. Fleming IA, Hindar K, MjÖlnerÖd IB, Jonsson B, Balstad T, Lamberg A (2000) Lifetime success and interactions of farm salmon invading a native population. Proceedings of the Royal Society of London B 267: 1517–1523.

16. Ford MJ (2002) Selection in captivity during supportive breeding may reduce fitness in the wild. Conservation Biology 16, 815–825.

17. Fushimi H (2001) Production of juvenile marine finfish for stock enhancement in Japan. Aquaculture 200 (1–2): 33–53.

18. Goshima S, Fujiwara H (1994) Distribution and abundance of cultured scallop *Patinopecten yessoensis* in extensive sea beds as assessed by underwater camera. Marine Ecology Progress Series 110: 151–158.

19. Hamasaki K (2008) Environmental change and resource enhancement: a case of abalone. In: Kitada S, Kaeriyama M, Hamasaki K, Taniguchi N (eds) Enhancement and Conservation of Fisheries Resources, pp. 86–104. Seizando. Tokyo [in Japanese].

20. Hamasaki K, Kitada S (2006) A review of kuruma prawn (*Penaeus japonicus*) stock enhancement programs in Japan. Fisheries Research 80: 80–90.

21. Hamasaki K, Kitada S (2008a) The enhancement of abalone stocks: lessons from Japanese case studies. Fish and Fisheries 9: 243–260.

22. Hamasaki K, Kitada S (2008b) Potential of stock enhancement for decapod crustaceans. Reviews in Fisheries Science 16: 164–174.

23. Hamasaki K, Kitada S (2013). Catch fluctuation of Kuruma prawn, *Penaeus japonicus* in Japan relative to ocean climate variability and a stock enhancement program. Reviews in Fisheries Science 21(3-4): 454–468.

24. Hamasaki K, Obata Y, Dan S, Kitada S (2011) A review of seed production and stock enhancement for commercially important portunid crabs in Japan. Aquaculture International 19: 217–235.

25. Hauser L, Carvalho GR (2008) Paradigm shifts in marine fisheries genetics: ugly hypotheses slain by beautiful facts. Fish and Fisheries 9: 333–362.

26. Hayakawa J, Yamakawa T, Aoki I (2007) Long-term fluctuation in the abundance of abalone and top shell in Japan and factors affecting those fluctuations. Bulletin of the Japanese Society of Fisheries Oceanography 71: 96–105 [In Japanese with English abstract].

27. Hilborn R (1992) Hatcheries and the future of salmon in the Northwest. Fisheries 17: 5–8.

28. Hilborn R (1998) The economic performance of marine stock enhancement projects. Bulletin of Marine Science 62: 661–674.

29. Hilborn R (2004) Population management in stock enhancement and sea ranching. In: Leber K, Kitada S, Blankenship HL, Svåsand T (eds) Stock Enhancement and Sea Ranching, Developments Pitfalls and Opportunities, pp. 201–210. Blackwell, Oxford.

30. Hokkaido Salmon Propagation Association (2017) Fifty years of Hokkaido Salmon Propagation Association. Sapporo [In Japanese].

31. Iida M, Yoshino K, Katayama S (2018) Current status of natural spawning of chum salmon *Oncorhynchus keta* in rivers with or without hatchery stocking on the Japan Seaside of northern Honshu, Japan. Fisheries Science 84: 453–459.

32. Ishino K (1999) Stocking effectiveness of Japanese flounder, *Paralichthys olivaceus*, fingerlings released on the south-western coast of Hokkaido Prefecture, Japan. In: Howel BR, Moksness E, Svåsand T (eds) Stock Enhancement and Sea Ranching, pp. 557–572. Blackwell, Oxford.

33. Kaeriyama M (1999) Hatchery programmes and stock management of salmonid populations in Japan. In: Howel BR, Moksness E, Svåsand T (eds) Stock Enhancement and Sea Ranching, pp. 153–167. Blackwell, Oxford.

34. Kaeriyama M, Seo H, Kudo H, Nagata M (2012) Perspectives on wild and hatchery salmon interactions at sea, potential climate effects on Japanese chum salmon, and the need for sustainable salmon fishery management reform in Japan. Environmental Biology of Fishes 94: 165–177.

35. Kaeriyama M, Seo H, Qin YX (2014) Effect of global warming on the life history and population dynamics of Japanese chum salmon. Fisheries Science 80: 251–260.

36. Kaeriyama M, Qin YX (2014) Biological interactions between wild and hatchery-produced Pacific salmon. In Woo PTK, Noakes DJ (eds), Salmon, pp. 223–238. Nova Science Publishers, New York.

37. Kasugai K, Saneyoshi H, Aoyama T, Shinriki Y, Iijima A, Miyakoshi Y (2016) Early marine migration of juvenile chum salmon along the Pacific coast of eastern Hokkaido. North Pacific Anadromous Fish Commssion Bulletin 6: 61–72.

38. Kayano Y, Tanaka T, Hayashi H (1998) Observations of artificially releasing seedlings and native fishes around the marine ranching area. Fisheries Engineering 35: 303–309 [In Japanese with English abstract].

39. Kitada S (1999) Effectiveness of Japan’s stock enhancement programs: current perspectives. In: Howel BR, Moksness E, Svåsand T (eds) Stock Enhancement and Sea Ranching, pp. 103–131. Blackwell, Oxford.

40. Kitada S (2001) Stock enhancement assessment with Japan examples. Kyoritsu, Tokyo, Japan [in Japanese].

41. Kitada S (2014) Japanese chum salmon stock enhancement: current perspective and future challenges. Fisheries Science 80: 237–249.

42. Kitada S (2018) Economic, ecological and genetic impacts of marine stock enhancement and sea ranching: A systematic review. Fish and Fisheries 19: 511–532.

43. Kitada S, Fujishima H (1997) The stocking effectiveness of scallop in Hokkaido. Nippon Suisan Gakkaishi 63: 686–693 [In Japanese with English abstract].

44. Kitada S, Hiramatsu K, Kishino H (1994) Estimating mortality rates from tag recoveries: incorporating over-dispersion, correlation, and change points. ICES Journal of Marine Science 51: 241–251.

45. Kitada S, Hirano K (1987) Estimation of mortality coefficients from tag recovery. Nippon Suisan Gakkaishi 53: 1765–1770 [In Japanese with English abstract].

46. Kitada S, Kishino H (2006) Lessons learned from Japanese marine finfish stock enhancement programmes. Fisheries Research 80: 101–112.

47. Kitada S, Nakajima K, Hamasaki K, Shishidou H, Waples RS, Kishino H (2019) Rigorous monitoring of a large-scale marine stock enhancement program demonstrates the need for comprehensive management of fisheries and nursery habitat. Scientific Reports 9: 5290.

48. Kitada S, Nakamichi R, Kishino H (2017) The empirical Bayes estimators of fine-scale population structure in high gene flow species. Molecular Ecology Resources 17: 1210–1222.

49. Kitada S, Okouchi H (1994) A simulation model for strategies of hatchery releases with application to red sea bream. Nippon Suisan Gakkaishi 60: 235–240 [In Japanese with English abstract].

50. Kitada S, Shishidou H, Sugaya T, Kitakado T, Hamasaki K, Kishino H (2009) Genetic effects of long-term stock enhancement programs. Aquaculture 290: 69–79.

51. Kitada S, Taga Y, Kishino H (1992) Effectiveness of a stock enhancement program evaluated by a two-stage sampling survey of commercial landings. Canadian Journal of Fisheries and Aquatic Sciences 49: 1573–1582.

52. Kitada S, Yoshikai R, Fujita T, Hamasaki K, Nakamichi R, Kishino H (2017) Population structure and persistence of Pacific herring following the Great Tohoku earthquake. Conservation Genetics 18: 423–437.

53. Kojima H (1995) Evaluation of abalone stock enhancement through the release of hatchery-reared seeds. Marine and Freshwater Research 46: 689–695.

54. Kondo H, Hirano Y, Nakayama N, Miyake M (1965) Offshore distribution and migration of Pacific salmon (genus *Oncorhynchus*) based on tagging studies (1958-1961). Bulletin of International North Pacific Fisheries Commission 17: 1–213.

55. Koya Y (2005) Stock enhancement of barfin flounder in Hokkaido. Technical Reports of Hokkaido Fisheries Experimental Station 5: 43–49 [In Japanese].

56. Kudo K, Kimoto H (1994) Fishery management of marine ranching in the Oita prefectural sea area. Fisheries Engineering 31:121–126 [In Japanese with English abstract].

57. Kurata M (1999) On the decline in the growth of maricultured scallop, Patinopecten yessoensis, in the Okhotsk coastal area of Hokkaido. Research Reports of Hokkaido Fisheries Experimental Station 54: 25–32 [In Japanese].

58. Laikre L, Schwartz MK, Waples RS, Ryman N, GeM working group (2010) Compromising genetic diversity in the wild: Unmonitored large-scale release of plant and animals. Trends in Ecology and Evolution 25: 520–529.

59. Le Vay L, Carvalho GR, Quinitio ET, Lebata JH, Ut VN, Fushimi H (2007) Quality of hatchery-reared juveniles for marine fisheries stock enhancement. Aquaculture 268 (1-4): 169–180.

60. Lee SI, Zhang CI (2018) Evaluation of the effect of marine ranching activities on the Tongyeong marine ecosystem. Ocean Science Journal 53: 557–582.

61. Lee S G, Rahman MA (2018) Ecological stock enhancement programs (ESEPs) based fisheries rebuilding plan (FRP) in Korea. Journal of Environmental Biology 39: 936–942.

62. Leber K, Kitada S, Blankenship H, Svåsand T (Eds.) (2004) Stock enhancement and sea ranching: Developments, pitfalls and opportunities. Blackwell, Oxford, UK.

63. Masuda R, Tsukamoto K (1998) Stock enhancement in Japan: review and perspective. Bulletin of Marine Science 62: 337–358.

64. Matsuishi T, Kishino H, Numachi K (1995) A model of gene displacement by stocking activities. Nippon Suisan Gakkaishi 61: 326–330 [In Japanese with English abstract].

65. Matsumura Y (2005) Optimal release strategy of hatchery-produced ocellate puffer *Takifugu rubripe* in Ariake Sound by mark-recapture experiments, based on the stocking effectiveness for young-of-the-year. Nippon Suisan Gakkaishi 71: 805–814 [In Japanese with English abstract].

66. Matsumura Y (2006) Stocking effectiveness of hatchery-produced juveniles of ocellate puffer *Takifugu rubripe* in natal spawning ground of Ariake Sound. Nippon Suisan Gakkaishi 72: 1029–1038 [In Japanese with English abstract].

67. McGinnity P, Prodöhl P, Ferguson A, Hynes R, ó Maoiléidigh N, Baker N, et al. (2003) Fitness reduction and potential extinction of wild populations of Atlantic salmon, Salmo salar, as a result of interactions with escaped farm salmon. Proceedings of the Royal Society of London B 270: 2443–2450.

68. Ministry of Agriculture, Forestry and Fisheries (MAFF) (1964–2018) Annual statistics of fisheries and aquaculture production in 1966–2019. Association of Agriculture and Forestry Statistics, Tokyo [in Japanese].

69. Miyakoshi Y, Nagata M, Sugiwaka KI, Kitada S (2001) Commercial harvest of hatchery-reared masu salmon *Oncorhynchus masou* estimated by a coast-wide sampling program in Hokkaido, northern Japan, and the two-stage sampling schemes of landings. Fisheries Science 67: 126–133.

70. Miyakoshi Y, Urabe H, Saneyoshi H, Aoyama T, Sakamoto H, Ando D et al. (2012) The occurrence and run timing of naturally spawning chum salmon in northern Japan. Environmental Biology of Fishes 94: 197–206.

71. Miyakoshi Y, Nagata M, Kitada S, Kaeriyama M (2013) Historical and current hatchery programs and management of chum salmon in Hokkaido, northern Japan. Reviews in Fisheries Science 21: 469–479.

72. Moksness E, Støle R (1997) Larviculture of marine fish for sea ranching purposes: is it profitable? Aquaculture 155: 341–353.

73. Moksness E, Støle R, van der Meeren G (1998) Profitability analysis of sea ranching with Atlantic salmon (*Salmo salar*), Arctic charr (*Salvelinus alpinus*), and European lobster (*Homarus gammarus*) in Norway. Bulletin of Marine Science 62: 689–699.

74. Morita K (2014) Japanese wild salmon research: toward a reconciliation between hatchery and wild salmon management. NPAFC Newsletter 35: 4–14.

75. Morita K (2016) Trends in Russian chum salmon populations. FRA Salmonid Research Report 10, 38–40 [In Japanese].

76. Morita K, Saito T, Miyakoshi Y, Fukuwaka MA, Nagasawa T, Kaeriyama M (2006) A review of Pacific salmon hatchery programmes on Hokkaido Island, Japan. ICES Journal of Marine Science 63: 1353–1363.

77. Morita K, Takahashi S, Ohkuma K, Nagasawa T (2013) Estimation of the proportion of wild chum salmon *Oncorhynchus keta* in Japanese hatchery rivers. Nippon Suisan Gakkaishi 79, 206–213 [In Japanese with English abstract].

78. Murakami O (2012) Effect of large-scale release of barfin flounder in the western Pacific Hokkaido. Reports of Hokkaido Fisheries Experimental Station 706 [In Japanese].

79. Nagata M, Miyakoshi Y, Urabe H, Fujiwara M, Sasaki Y, Kasugai K et al. (2012) An overview of salmon enhancement and the need to manage and monitor natural spawning in Hokkaido, Japan. Environmental Biology of Fishes 94: 311–323.

80. Naish KA, Taylor III JE, Levin PS, Quinn TP, Winton JR, Huppert D et al. (2007) An evaluation of the effects of conservation and fishery enhancement hatcheries on wild populations of salmon. Advances in Marine Biology 53: 61–194.

81. Nakagawa M, Okouchi H, Adachi J (2004) Stocking effectiveness of black rockfish *Sebastes schlegeli* released in Yamada Bay evaluated by a fish market census. In: Leber K, Kitada S, Blankenship HL, Svåsand T (eds) Stock Enhancement and Sea Ranching, Developments Pitfalls and Opportunities, pp. 501–511. Blackwell, Oxford.

82. Nakajima H, Kai M, Koizumi K, Tanaka T., Machida M (2008) Optimal release locations of juvenile ocellate puffer *Takifugu rubripes* identified by tag and release experiments. Reviews in Fisheries Science: 16(1-3), 228–234.

83. Nakajima K, Kitada S, Yamazaki H, Takemori H, Obata Y, Iwamoto A et al. (2013) Ecological interactions between hatchery and wild fish: a case study based on the highly piscivorous Japanese Spanish mackerel. Aquaculture Environment Interactions 3: 231–243.

84. Nakajima K, Kitada S, Habara Y, Sano S, Yokoyama E, Sugaya T et al. (2014) Genetic effects of marine stock enhancement: a case study based on the highly piscivorous Japanese Spanish mackerel. Canadian Journal of Fisheries and Aquatic Sciences 71: 301–314.

85. Nakamura A, Kitada S, Hamasaki K, Okouchi H (2005) Catch fluctuation in *Haliotis* spp. abalones: annual landings of the Ezo abalone *Haliotis discus hannai* in relation to climate oscillation and stock enhancement programs. Saibai Gyogyo Gijyutsu Kaihatsu Kenkyu 33: 45–54 [In Japanese with English abstract].

86. Nei M, Chesser RK (1983) Estimation of fixation indices and gene diversities. Annals of Human Genetics 47: 253–259.

87. Nishihama Y (2001) Scallop fisheries in the Okhotsk Sea. Hokkaido University Press, Sapporo [in Japanese].

88. North Pacific Anadromous Fish Commission (NPAFC) (2019) NPAFC Pacific salmonid catch and hatchery release data (updated 2019 summer). NPAFC, Vancouver, BC. Retrieved from http://www.npafc.org

89. North Pacific Japan Stock Ehancement Counci (NPJSEC) (2015) Barfin flounder enhancement plan 2015–2021. Fisheries Agency [In Japanese].

90. Obata Y, Imai H, Kitakado T, Hamasaki K, Kitada S (2006) The contribution of stocked mud crabs *Scylla paramamosain* to commercial catches in Japan, estimated using a genetic stock identification technique. Fisheries Research 80: 113–121.

91. Obata Y, Iwamoto A, Takemori H, Yamazaki H, Okumura S, Fujimoto H et al. (2007) A comparison of survival rates until recruitment for hatchery-released Japanese Spanish mackerel *Scomberomorus niphonius* with different sizes at release. Nippon Suisan Gakkaishi 73: 55–61 [In Japanese with English abstract].

92. Obata Y, Yamazaki H, Iwamoto A, Hamasaki K, Kitada S (2008) Evaluation of stocking effectiveness of the Japanese Spanish mackerel in the eastern Seto Inland Sea, Japan. Reviews in Fisheries Science 16: 35−24.

93. Ohnuki T, Morita K, Tokuda H, Okamoto Y, Ohkuma K (2015) Numerical and economic contributions of wild and hatchery pink salmon to commercial catches in Japan estimated from mass otolith markings. North American Journal of Fisheries Management 35: 598–604.

94. Okamoto K (2004) Juvenile release and market size recapture of the swimming crab *Portunus trituberculatus* (Miers) marked with coded wire tags. In: Leber K, Kitada S, Blankenship HL, Svåsand T (eds) Stock Enhancement and Sea Ranching, Developments Pitfalls and Opportunities, pp. 181–186. Blackwell, Oxford.

95. Okouchi H, Kitada S, Tsuzaki T, Fukunaga T, Iwamoto A (1999) Number of returnes and economic return rates of hatchery-released flounder *Paralichthys olivaceus* in Miyako Bay – Evaluation by a fish market census. In: Howel BR, Moksness E, Svåsand T (eds) Stock Enhancement and Sea Ranching, pp. 573–582. Blackwell, Oxford.

96. Okouchi H, Kitada S, Iwamoto A, Fukunaga T (2004) The flounder stock enhancement in in Miyako Bay. FAO Fisheries Technical Paper 429: 171–202.

97. Okouchi H, Kitada S, Morioka T, Imamura S (1994) A comparison of the quality of hatchery-reared red sea bream *Pagrus major* based on mortality rate estimated from tag recoveries. Nippon Suisan Gakkaishi 60: 229–233 [in Japanese with English abstract].

98. Quinn TP (1993) A review of homing and straying of wild and hatchery-produced salmon. Fisheries Research 18(1-2): 29–44.

99. Reisenbichler RR, McIntyre JD (1977) Genetic differences in growth and survival of juvenile hatchery and wild steelhead trout, Salmo gairdneri. Journal of the Fisheries Board of Canada 34: 123–128.

100. Roberge C, Normandeau E, Einum S, Guderley H, Bernatchez L (2008) Genetic consequences of interbreeding between farmed and wild Atlantic salmon: insights from the transcriptome. Molecular Ecology 17: 314–324.

101. Sakai Y, Tajima K, Agatsuma Y (2004) Stock enhancement of the short-spined sea urchin. In: Leber K, Kitada S, Blankenship HL, Svåsand T (eds) Stock Enhancement and Sea Ranching, Developments Pitfalls and Opportunities, pp. 465–476. Blackwell, Oxford.

102. Sakano E (1960) Results from marking experiments on young chum salmon in Hokkaido, 1951-1959. Scientific Reports of the Hokkaido Salmon Hatchery 15: 17–38 [in Japanese with English Abstract].

103. Sato S, Templin WD, Seeb LW, Seeb JE, Urawa S (2014) Genetic structure and diversity of Japanese chum salmon populations inferred from single-nucleotide polymorphism markers. Transactions of the American Fisheries Society 143: 1231–1246.

104. Shiota K, Kitada S (1992) Life history of swimming crab in Hiuchi Nada, Seto Inland Sea estimated from marking experiments. Nippon Suisan Gakkaishi 58: 2297–2302 [in Japanese with English abstract].

105. Shishidou H (2002) Stocking effectiveness of red sea bream *Pagrus major* in Kagoshima Bay. Fisheries Science 68 (Suppl I): 904–907.

106. Shishidou H, Kitada S (2007) Stocking effectiveness of red sea bream *Pagrus major* in Kagoshima Bay, Japan. Nippon Suisan Gakkaishi 73: 270–277 [in Japanese with English abstract].

107. Shishidou H, Takimoto A, Obata Y, Hamasaki K, Kitada S (2012) Stock assessment and release strategy of red sea bream Pagrus major in Kagoshima Bay, Japan. Nippon Suisan Gakkaishi 78: 161–170 [in Japanese with English abstract].

108. Sproul JT, Tominaga O (1992) An economic review of the Japanese flounder stock enhancement project in Ishikari Bay, Hokkaido. Bulletin of Marine Science 50: 75–88.

109. Steward CR, Bjornn TC (1990) Supplementation of salmon and steelhead stocks with hatchery fish: A synthesis of published literature. *U.S. Fish and Wildlife Services*, Technical Reports 90 (1), 202 p.

110. Svåsand T, Kristiansen TS, Pedersen T, Salvanes AV, Engelsen R, Naevdal G et al. (2000) The enhancement of cod stocks. Fish and Fisheries 1: 173–205.

111. Takaba M, Kitada S, Nakano H, Morioka T (1995) Survival, feeding activity and liver constituents of red sea bream released into the Seto Inland Sea of Japan with special reference to fish quality. Nippon Suisan Gakkaishi 61: 574–579 [in Japanese with English abstract].

112. Takami H, Saido T, Endo T, Noro T, Musashi T, Kawamura T (2008) Overwinter mortality of young-of-the-year Ezo abalone in relation to seawater temperature on the North Pacific coast of Japan. Marine Ecology Progress Series 367: 203–212.

113. Takeuchi T (2001) A review of feed development for early life stages of marine finfish in Japan. Aquaculture 200(1–2): 203–222.

114. Taniguchi N (2003) Genetic factors in broodstock management for seed production. Reviews in Fish Biology and Fisheries 13: 177–185.

115. Taniguchi N (2004) Broodstock management for stock enhancement programs of marine fish with assistance of DNA markers (a review). In: Leber K, Kitada S, Blankenship HL, Svåsand T (eds) Stock Enhancement and Sea Ranching, Developments Pitfalls and Opportunities, pp. 329–338. Blackwell, Oxford.

116. Taylor MD, Chick RC, Lorenzen K, Agnalt AL, Leber KM, Blankenship HL et al. (2017) Fisheries enhancement and restoration in a changing world. Fisheries Research 186: 407–412.

117. Tillotson MD, Barnett HK, Bhuthimethee M, Koehler ME, Quinn TP (2019) Artificial selection on reproductive timing in hatchery salmon drives a phenological shift and potential maladaptation to climate change. Evolutionary Applications 12: 1344–1359.

118. Tominaga O, Watanabe Y (1998) Geographical dispersal and optimum release size of hatchery-reared Japanese flounder *Paralichthys olivaceus* released in Ishikari Bay, Hokkaido, Japan. Journal of Sea Research 40(1–2): 73–81.

119. Tomiyama T, Watanabe M, Fujita T (2008) Community-based stock enhancement and fisheries management of the Japanese flounder in Fukushima, Japan. Reviews in Fisheries Science 16(1–3): 146–153.

120. Tsujimura H (2007) Stocking effectiveness of red spotted grouper in Osaka Bay. www.kannousuiken-osaka.or.jp [In Japanese].

121. Tsukamoto K, Kuwada H, Uchida K, Masuda R, Sakakura A (1999) Fish quality and stocking effectiveness: behavioural approach. In: Howel BR, Moksness E, Svåsand T (eds) Stock Enhancement and Sea Ranching, pp. 203–218. Blackwell, Oxford.

122. Uki N (2006) Stock enhancement of the Japanese scallop *Patinopecten yessoensis* in Hokkaido. Fisheries Research 80: 62–66.

123. Wada T, Kamiyama K, Shimamura S, Mizuno T, Nemoto Y (2012) Effectiveness of stock enhancement of a rare species, spotted halibut *Verasper variegatus*, in Fukushima, Japan. Aquaculture 364: 230–239.

124. Wang Y, Yu J, Chen P (2018) Remote sensing assessment of ecological effects of marine ranching in the eastern Guangdong waters, China. Journal of Geoscience and Environment Protection 6: 101–113.

125. Waples RS (1991) Genetic interactions between hatchery and wild salmonids: lessons from the Pacific Northwest. Canadian Journal of Fisheries and Aquatic Sciences 48 (S1), 124–133.

126. Waples RS (1999) Dispelling some myths about hatcheries. Fisheries 24: 12–21.

127. Waples RS, Drake J (2004) Risk/benefit considerations for marine stock enhancement: A Pacific salmon perspective. In: Leber K, Kitada S, Blankenship HL, Svåsand T (eds) Stock Enhancement and Sea Ranching, Developments Pitfalls and Opportunities, pp. 260–306. Blackwell, Oxford.

128. Waples RS, Gaggiotti O (2006) What is a population? An empirical evaluation of some genetic methods for identifying the number of gene pools and their degree of connectivity. Molecular Ecology 15: 1419–1439.

129. Winton J, Hilborn R (1994) Lessons from supplementation of chinook salmon in British Columbia. North American Journal of Fisheries Management 14: 1–13.

130. Yamaguchi T, Ito S, Hamasaki K, Kitada S (2006) Stocking effectiveness of hatchery-released kuruma prawns estimated by two-stage sampling of commercial catch in Ariake Sound, Japan. Fisheries Science 72: 233–238.

131. Zhou X, Zhao X, Zhang S, Lin J (2019) Marine ranching construction and management in East China Sea: Programs for sustainable fishery and aquaculture. Water 11: 1237.

